# A retrospective analysis of 400 publications reveals patterns of irreplicability across an entire life sciences research field

**DOI:** 10.1101/2025.07.07.663460

**Authors:** Joseph Lemaitre, Désirée Popelka, Blandine Ribotta, Hannah Westlake, Sveta Chakrabarti, Li Xiaoxue, Mark A. Hanson, Haobo Jiang, Francesca Di Cara, Estee Kurant, Fabrice David, Bruno Lemaitre

**Author notes:** Corresponding authors: Joseph Lemaitre, Bruno Lemaitre.

## Abstract

The *ReproSci* project retrospectively analyzed the conceptual replicability of 1006 claims from 400 papers published between 1959 and 2011 in the field of *Drosophila* immunity. This project represents an expert assessment of the replicability 14 years later, of the replicability of nearly all publications across an entire scientific community in experimental life sciences. We found that 61% of claims were verified, while only 7% were directly challenged (not replicable), a replicability rate higher than previous assessments. Notably, 24% of claims had never been independently tested and remain unchallenged. We performed experimental validations of a selection of 45 unchallenged claim, that revealed that a significant fraction (38/45) of them is in fact non-replicable. We also found that claims from top-ranked institutions were more likely to be challenged and that claims published in trophy journals showed a similar but weaker trend. In line with the reproducibility crisis narrative, the rates of both challenged and unchallenged claims increased over time, especially as the field gained popularity. We characterized the uneven distribution of irreplicability among first and last authors. Surprisingly, irreplicability rates were similar between PhD students and postdocs, and did not decrease with experience or publication count. However, group leaders, who had prior experience as first authors in another *Drosophila* immunity team, had lower irreplicability rates, underscoring the importance of early-career training. Finally, authors with a more exploratory, short-term engagement with the field exhibited slightly higher rates of challenged claims and a markedly higher proportion of unchallenged ones. While limited by its reliance on published literature, this systematic, field-wide retrospective study offers meaningful perspective into the ongoing discussion on reproducibility in experimental life sciences.

## Introduction

Science predominantly thrives on the continuous aggregation of knowledge, with researchers progressively building upon one another’s discoveries. As results are replicated (when independent studies reach the same conclusions as the original authors of a claim), confidence in the evidence supporting the claim grows. Confidence in publishing results is critical to sustain trust within a scientific community and to allow new research to build on previous findings (Fidler and Wilcox, 2018). Over the past decades, this confidence has been on a decline.

Several recent reports point out that replicability is lower than desirable in molecular life sciences ( The Brazilian Reproducibility Initiativel 2026; Baker, 2016; DasGupta et al., 2007; Errington et al., 2021b; Prinz et al., 2011). In 2016, an extensive survey indicated that more than 70% of researchers have tried and failed to reproduce another scientist’s experiments, and more than half have failed to reproduce their own experiments (Baker, 2016). Another study suggested that about 85% of biomedical research is ‘wasted’, meaning that inefficiencies and methodological flaws throughout the research process undermine the value of scientific findings (Begley and Ioannidis, 2015; Chalmers and Glasziou, 2009). In addition, 52% of scientists that were surveyed by Nature agree that we are currently facing a significant ‘crisis’ of replicability. Ongoing concerns about replicability in academic research carry important consequences for the private sector. Two pharmaceutical companies, Bayer and Amgen, reported replicability rates of 11% and 25% in two independent efforts to reproduce findings from dozens of groundbreaking basic studies in cancer and related areas (Prinz et al., 2011). Moreover, an extensive project in cancer biology reveals that a very significant fraction of findings reported in high profile journals cannot be replicated or have smaller effect size that initially reported (Errington et al., 2021, 2014). Lack of replicability can have multiple causes, ranging from the effect of cryptic factors not accounted for by authors, misuse of technical tools, spoiled reagents, bias in data reporting, lack of statistical power or even outright fraud (Devezer and Buzbas, 2023; Eisner, 2018; Lesperance and Broderick, 2021; Peng, 2015).

Misleading or erroneous findings can disrupt scientific progress, wasting valuable time and resources as researchers attempt to replicate or use the results of a published article (Stern et al., 2014). In some cases, entire research initiatives may be built upon findings that are later discovered to be inaccurate or flawed (Piller, 2022). This might also result in unfair career outcomes if sensational but irreproducible results are rewarded over rigorous studies, especially in high-impact publications (Lemaitre, 2016; Steneck, 2011). However, full replicability is neither realistic nor necessarily desirable. A certain level of irreplicability may reflect scientific boldness and risk-taking. Cutting-edge explorations of the unknown are expected to involve occasional backtracking as the bigger picture gradually comes into focus. Thus, striving for perfect replicability could paradoxically stifle innovation and promote overly conservative science. Determining the ideal balance is currently an open question (Shiffrin et al., 2018).

Considering the prominent role played by science in contemporary society, there is a growing need to understand how academic incentives influence the reliability of published results. Indeed, the field of metascience - the scientific investigation of science itself - is gaining popularity (Fortunato et al., 2018). Analyzing how replicability affects the scientific enterprise is required to better characterize how science progresses despite mistakes. It can also help to influence policy to improve science replicability, or at least to limit its most detrimental impacts.

Research focusing on the immune system of *Drosophila* flourished in the early 1990s and is still an active field of investigation. It boasts two thousand articles published since its beginning, dealing with various facets of the immune response of this insect, such as humoral immunity, cellular immunity, response to pathogens, immuno-metabolism, and evolution of the immune system (Westlake et al., 2024). While the field remains modest in size, it gained visibility in the mid 1990s when researchers realized that some mechanisms regulating the *Drosophila* immune defense are conserved in mammals, including humans. The discovery of Toll-like receptors, stemming from the identification of Toll’s role in the *Drosophila* immune response (Lemaitre, 2004), established the value and utility of the field to immunological research, a finding that was acknowledged by a Nobel prize awarded to Jules Hoffmann in 2011.

The medium size of this research field and the lack of direct biomedical influence are ideal for a metascience analysis. The *Drosophila* immunity field is substantially smaller than other fields of research such as HIV, microbiota or cancer, where the rapid accumulation of articles forbids any comprehensive evaluation of scientific claims by an individual or a laboratory. With fewer direct translational or biomedical influences, we expect scientists working on *Drosophila* immunity to be less affected by external biases, and to prioritize recognition by peers. Moreover, *Drosophila* research is generally perceived as solid, because it relies predominantly on its powerful genetics and well-controlled experimental conditions, so the field is particularly interesting for this analysis.

In the framework of the *ReproSci* project, we retrospectively analyzed the replication of 400 articles published between 1959 and 2011 in the field of *Drosophila* immunity research-that is, nearly all the major articles published by a community of scientists in experimental life science over 50 years (Westlake et al., 2026). This expert assessment project focuses on assessing the strength of the claims themselves (i.e. “indirect reproducibility” or “conceptual replicability”) rather than testing whether the original methods produce repeatable results (“direct reproducibility”). Thus, our conclusions do not directly challenge the initial results leading to a claim, but rather the general applicability (i.e. generalizability) of the claim itself (Goodman et al., 2016; Meng, 2020). For each article, we identified the main, major, and minor scientific conclusions, hereafter referred to as “claims”. The 14-year interval between the time the latest article was published and our study, during which scientific knowledge has advanced and other laboratories have had the opportunity to attempt replication, provides the necessary perspective to efficiently assess the validity of the claims. In a first step, we created a publicly accessible website, displaying all the information on the articles, the claims, and their validity (https://ReproSci.epfl.ch/). Second, we contacted the *Drosophila* scientific community to provide feedback via this website on the replicability of the claims. In a third step, several teams attempted to replicate 45 major and 11 minor claims that remained unchallenged in the literature at the time. The present article focuses on the meta-science findings, while the scientific consequences of the *ReproSci* project in the *Drosophila* immunity field are presented in a companion article (Westlake et al., 2026).

This article analyzes the pattern of replicability with two objectives: first, to comprehend the level of replicability within the experimental life science community, and second, to examine whether instances of non-replicability tend to aggregate among journals, first authors, and lead researchers. Such clustering may identify individual characteristics and environmental factors that influence science replicability. We open the Results section with a detailed descriptive analysis of the effect of individual categorical variables on reproducibility outcomes. Then, to assess the joint influence of all covariates and account for the clustering of multiple claims by the same author, we fit a multivariate model to account for confounding effects across our covariates. Because challenged claims are rare and the overall sample is modest, statistical power remains limited. However, taken together, i) the descriptive and ii) multivariate analyses form a single narrative, the former describing the observed patterns exhaustively, and the latter testing the robustness of the associations.

## Results

### Methodology and claim assessment

A list of 400 publications published before 2011 was generated using a curated search string on the publicly available PubMed database (Westlake et al., 2026). This limitation was imposed to allow a time frame in which follow-up studies may be published that may verify, expand on or contradict the content of the publications selected for the study. Scientific claims were divided into three categories. The main (one per article), major (3-4) and minor (2-4) claims were manually extracted from 400 articles in the field of *Drosophila* immunity published before 2011. We then assessed the claims of each article to determine if they were supported or refuted by subsequent publications. The claims were manually categorized by H.W. and B.L. into four broad groups:

i. verified by subsequent studies when later studies confirmed the claim,
ii. challenged when later studies contradicted or disproved the claim,
iii. unchallenged when no later study addressed or attempted to replicate the claim,
iv. partially verified or mixed when later studies did not lead to a definitive conclusion

Additional description of the categorization can be founbd in **Table 1** in Methods for classification, the companion article (Westlake et al., 2026), and on the *ReproSci* website (https://reprosci.epfl.ch/). Then, experimental work was performed in several laboratories to test the validity of 45 major and 11 minor unchallenged claims using alternate methods. These claims subjected to experimental validation were selected according to criteria such as appearing suspicious due to the absence of direct follow up or being straightforward to test experimentally. Our data and findings were put on a community-accessible website that opened to the public in early July 2023. The present article focusses on the pattern of irreplicability and statistical analyses were performed on the 1006 major claims only (referred to as claims below) of the database as updated on December 20^th^ 2024. In this article, challenged claims will interchangeably be called irreplicable.

**Table 1.** Verification categories used to assess claims.

| Category | Characteristics |
|---|---|
| *Verified* | • An independent lab or lab(s) have generated results that directly support this claim using similar or alternate methods. |
| *Verified by same authors* | • Subsequent work by the same lab or authors supports or extends this claim. |
| *Verified by reproducibility project* | • This claim is supported by experiments executed specifically to assess it in the Lemaitre Lab or other laboratories in collaboration with the *ReproSci* project. |
| *Partially verified* | • Part of the claim is verified by additional data, but part of it is contradicted<br>**OR**<br>• The claim is not supported by subsequent research, but the data used to justify the claim are consistent with current knowledge. The data were interpreted incorrectly either due to lack of context at the time or other confounding factors.<br>**OR**<br>• The claim is supported by subsequent research, but the data used to justify the claim in the original publication are not. |
| *Unchallenged* | • No additional data is available that significantly supports or contradicts this claim. |
| *Unchallenged, logically consistent* | • No additional data is available that directly supports this claim, but the claim is consistent with other well-supported related data in the field. |
| *Unchallenged, logically inconsistent* | • No additional data is available that directly contradicts this claim, but the claim is inconsistent with other well-supported related data in the field. |
| *Mixed* | • Both data that supports and contradicts this claim have been published in the field. It is unclear based on the published data whether the claim is correct, incorrect, or whether our understanding of the topic is simply incomplete. |
| *Challenged* | • An independent lab or lab(s) have generated results that directly contradict this claim using similar or alternate methods. |
| *Challenged by same authors* | • Subsequent work by the same lab or authors contradicts this claim. |
| *Challenged by reproducibility project* | • This claim is contradicted by experiments executed specifically to assess it in the Lemaitre Lab or other laboratories in collaboration with the *ReproSci* project.<br>• When complete, a link to a writeup of the experimental results is included. |

### *Drosophila* immunity claims are mostly replicable

After an extensive literature review to assess the validity of claims published from 1959 to 2011, combined with experimental testing of 45 major claims, we found that approximately 61% of major claims (610 out of 1006 across the 400 selected articles) were fully verified (**Table 2**). Among these 610 verified claims, 91.6% were corroborated by independent laboratories, 7.2% were confirmed by the same laboratory, and 1.1% were verified through experimental work conducted as part of the *ReproSci* project. This verification rate is higher than that observed in other fields where reproducibility has been systematically assessed ( The Brazilian Reproducibility Initiative 2026; Amaral et al., 2019; Da Costa et al., 2024; Errington et al., 2021; Fidler et al., 2017; Youyou et al., 2023). These results may reflect the robust scientific standards and methodological rigor of *Drosophila* research, which benefits from the availability of diverse and powerful genetics approaches and from a collaborative culture of rapid reagent sharing, that facilitates replication across laboratories.

**Table 2:** Proportion of claims in broad reproducibility assessment categories before and after experimental validation by the *ReproSci* project. See description of each category in Material and Methods. In this article, we are using indirect reproducibility, also called conceptual replicability, to assess the validity of a claim using alternate methods to those used in the original study.

| **Major claim** | **Nb (%)** | **Nb (%) after experimental validation** |
|---|---|---|
| Verified | 603 (59.9%) | 610 (60,6%) |
| Unchallenged | 285 (28,3%) | 240 (23,8%) |
| Partially verified | 75 (7,5 %) | 75 (7,5%) |
| Mixed | 12 (1,2%) | 12 (1,2%) |
| Challenged | 31 (3,1%) | 69 (6,8%) |
| Total | 1006 | 1006 |

We also observed that 23.8% of major claims had no published follow-up (**Table 2**, **Figure 1**); these unchallenged claims are particularly interesting and will be discussed in detail in the next section. Additionally, 7.5% of the claims were categorized as partially verified. This category encompasses cases where insightful data were accompanied by incomplete interpretations, or where incomplete data were paired with insightful interpretations. Importantly, 6.8% of claims (69 out of 1006) were challenged. Among these, 44.9% (31 out of 69) were contested by published articles including 7.2% by the same authors, and 55.1% (38 out of 69) were challenged experimentally as part of the *ReproSci* project. Thus, the experimental validation performed during this project doubled the number of reported challenged claims. Finally, 1.2% of major claims were classified as “mixed” (**Table 2**). This category includes claims that presented particularly challenging issues (e.g., conflicting data), making it impossible to definitively categorize them as either verified or challenged.

**Figure 1:**
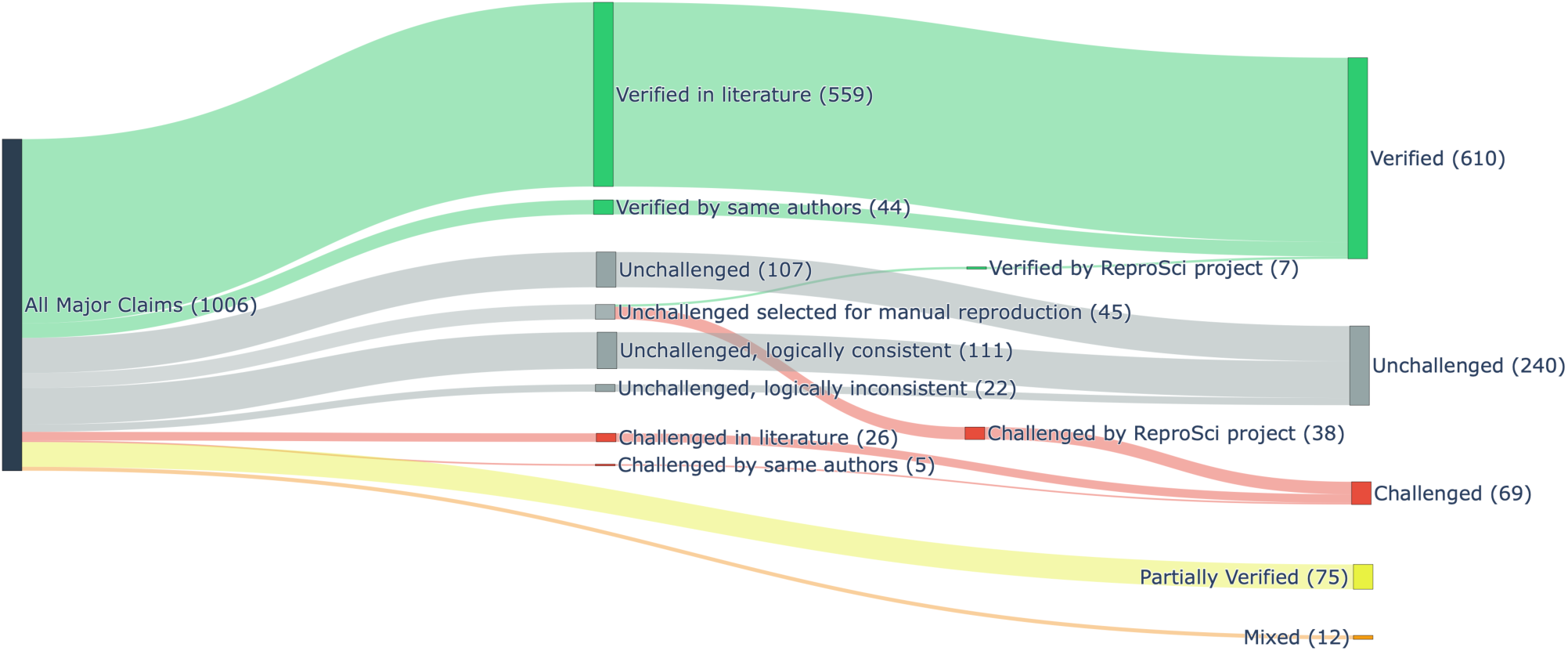
Flow of the study. Claims were first classified in five categories based on literature. 45 unchallenged claims were selected to be experimentally tested and classified either as challenged or as verified. We further classified some unchallenged as *consistent* or *inconsistent* depending on their consistency with current knowledge.

In conclusion, contrary to the prevailing “reproducibility crisis” narrative, a significant proportion of the findings published in the field of *Drosophila* immunity before 2011 has been subsequently verified by independent studies.

### A significant fraction of unchallenged claims is non-reproducible

Our analysis highlights a significant proportion of “unchallenged” claims that were not followed by subsequent studies and whose validity was unknown prior to our experimental evaluation. Most reproducibility studies have focused on verified or challenged claims so little is known about the vast number of statements generated in a scientific community without subsequent direct validation. Despite the lack of validation, these “unchallenged claims” are often assumed to be valid and contribute to the broader construction of scientific knowledge. An important challenge is to estimate the percentage of unchallenged major claims that are actually non-replicable.

Prior to our experimental validation, we identified a total of 285 unchallenged claims, representing 28.3% of all major claims published up to 2011. We conducted experimental evaluations of 45 unchallenged major claims with the assistance of multiple laboratories (Westlake et al., 2026). The selection of unchallenged claims for testing was intentionally biased toward two categories: (i) those that appeared suspicious in light of the existing literature (31 claims), and (ii) those that were relatively straightforward to test experimentally (14 claims). Among the 45 unchallenged major claims tested, 84% (38 out of 45) could not be experimentally replicated and were then reclassified as “challenged by the *ReproSci* project.” Conversely, only 16% (7 out of 45) were successfully replicated and reclassified as “verified by the *ReproSci* project” (**Figure 1**). Notably, all 31 suspicious unchallenged claims were challenged, while 7 out of the 14 unchallenged “easy to test” claims were also challenged. Excluding the 45 unchallenged major claims that were experimentally tested, we categorized the remaining 240 unchallenged claims into three groups. The first one referred to as ‘unchallenged logically consistent’ are claims not directly tested but consistent with current knowledge of the field (111 claims). The second referred to as ‘unchallenged logically inconsistent’ are claims that appear inconsistent with current knowledge (22 claims). Finally the third group, referred to as ‘unclassified’ are claims that could not be definitively categorized (107 claims) (**Figure 1**). In the remainder of the text, we analyze the claims following *ReproSci* experimental validation, labelling the 45 selected claims as either challenged or verified.

This analysis reveals that a substantial number of claims remain unchallenged even a decade after publication. Moreover, our findings suggest that a considerable proportion of these claims may not be replicable.

### Higher representation of challenged claims in trophy journals

Articles published in high-impact journals receive high visibility, a key factor in career progression and funding security (van Wesel, 2016). Additionally, high-impact publications are increasingly used as metrics to evaluate university rankings. Consequently, scientists face significant internal and external pressures to publish in such prestigious journals (Udesky, 2025). On the other hand, high-impact journals aim to attract the most surprising findings, which are inherently more likely to be challenged than more conventional studies. Based on this, we hypothesized that articles published in high-impact journals are more likely to include non-replicable results. To test this hypothesis, we estimated the number of challenged claims published in journals categorized as having low or medium-impact factors (<10), high-impact factors (>10 and <50), and “trophy journals” of the highest reputation (>50, which corresponds to *Science*, *Nature*, and *Cell* only) using the 2022 impact factors. The results are summarized in **Figure 2**, with a more detailed breakdown by journal provided in **Table S1**. Compared with low-impact journals, where 5.8 % of major claims were challenged, the proportion rose to 7.8 % in high-impact journals (odds ratio [OR] = 1.36, 95 % confidence interval [CI] 0.80–2.32) and more than doubled to 12.9 % in trophy journals (OR = 2.39, 95 % CI 1.06–5.40). We recall that the OR expresses how much more (OR > 1) or less (OR < 1) likely a claim is to be challenged relative to articles published in low-impact journals, while the 95 % CI indicates the range of values compatible with random sampling error; OR intervals that exclude 1 denote a statistically significant effect. Thus, our analysis shows that major claims published in low-impact journals are significantly more likely to be replicable than major claims published in trophy journals. One explanation for this trend is that claims published in high-impact journals are more frequently subjected to replication attempts or used as a foundation for further research, thereby increasing the likelihood of identifying irreplicable results. Indeed, low-impact journals include a higher share of claims remaining unchallenged (29.3 %, 95 % CI 25.9–32.9 %) compared to high-impact journals (13.9 %, 95 % CI 10.5–18.2 %) and to trophy journals (17.7 %, 95 % CI 10.2–29.0 %). However, variation in the number of unchallenged claims does not explain the higher proportion of challenged claims in trophy journals compared to high-impact journals. If we take into consideration the percentage of verified, unchallenged and challenged claims, **Figure 2B** shows that high-impact but not trophy or low-impact journals contain the highest proportion of verified claims. Thus, we conclude that while trophy journals may indeed attract some of the most impactful science, they also appear more likely to include irreplicable claims.

**Figure 2.**
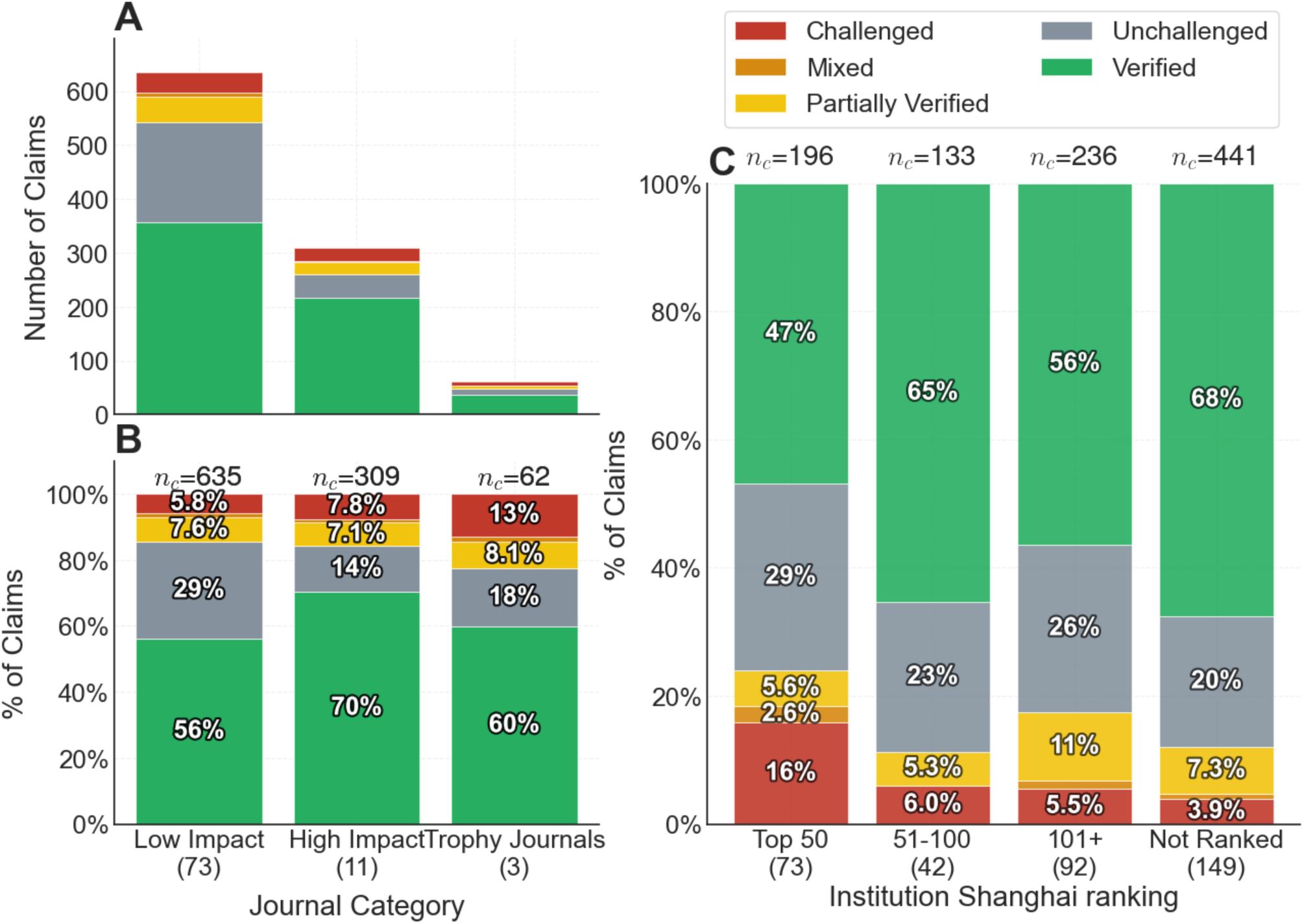
Percentage of claims by category according to journal-impact and institution ranking. (**A**, **B**) The distribution of claims across the five assessed categories for articles published in low-impact, high-impact, and trophy journals. The number of claims is indicated at the top. Panel A presents raw counts, while Panel B shows normalized data. **(C)** The distribution of claims across the five assessed categories according to the ranking of the last author institution in Shanghai Ranking’s 2010 Academic Ranking of World Universities. The number of claims is indicated at the top, while the number of institutions in each category is indicated at the bottom. The Not Ranked categories encompass research institutes (ex. CNRS), hospitals and other institution that are not included in the Shanghai ranking. In this figure, and in the rest of the paper, the number n_c_ indicates the number of claims in each category, while the number at the bottom highlight the count in each category (e.g there are 11 high-impact journals, and 73 Top-50 institutions).

### Higher representation of challenged claims from top universities

We then investigated whether the percentage of irreplicability varies among laboratories according to the prestige of their affiliate universities. To test this hypothesis, we classified all the major claims published according to the Shanghai ranking of the last author university. **Figure 2C** shows that claims from laboratories at top-50 universities were challenged far more often 15.8 % (95 % CI 11.4–21.6 %) than other institutions, up to a four-fold increase in the odds of being questioned (OR = 4.69, 95 % CI 2.53–8.70) versus 3.9 % for unranked institutions, which notably include hospitals and research institutes such as the French CNRS. Even after adjusting for journal tier, see multivariate model below, university prestige remains the dominant signal: claims from top-50 institutions are 3.6 times more likely to be challenged than those from unranked universities (OR 3.68, 94 % HDI 1.50–9.28). By comparison, publishing in a “trophy” journal adds only a notable and uncertain risk (OR 1.75 0.59–4.91), while other journal or ranking bands show no clear effect. Put simply, elite affiliation more than journal prestige best predicts irreplicability in our sample.

### The irreplicability rate has increased over time as the field has grown in popularity

The reproducibility crisis narrative suggests that irreplicability has increased over recent decades, posing a significant challenge to the life sciences (Baker, 2016; Hunter, 2017). Our dataset spans 50 years, from the onset of *Drosophila* immunity research as a relatively marginal field to its raise to prominence, with a notable increase in publications since the mid-1990s driven by growing interest in innate immunity. We analyzed fluctuations in irreplicability over time by grouping major claim assessments into time windows: pre-1991, followed by 5-year intervals until 2011. The percentage of irreplicability was calculated for each time window (**Figure 3**). Across the five windows examined, the proportion of challenged claims climbed steadily from 0% in the small corpus prior to 1991 (0/29 claims), to 4.6% during 1992–1996, and then further increased to 7.7% between 1997 and 2001. This upward trend peaked at 8.3% during the 2002–2006 period, before slightly declining to 6.1% between 2007 and 2011 (**Figure 3**). Despite these apparent temporal shifts, none of the pairwise comparisons between periods reached statistical significance, with p-values consistently ≥ 0.15. Thus, while the trends highlighted in our study are consistent with a raise of irreplicability, we cannot exclude that the observed upward trajectory might reflect sampling variation rather than a definitive sustained temporal trend. Further studies focusing on articles published after 2011 could clarify whether the mild decrease we observed marks the beginning of a downward trend or reflects fluctuations at a consistently high level.

**Figure 3.**
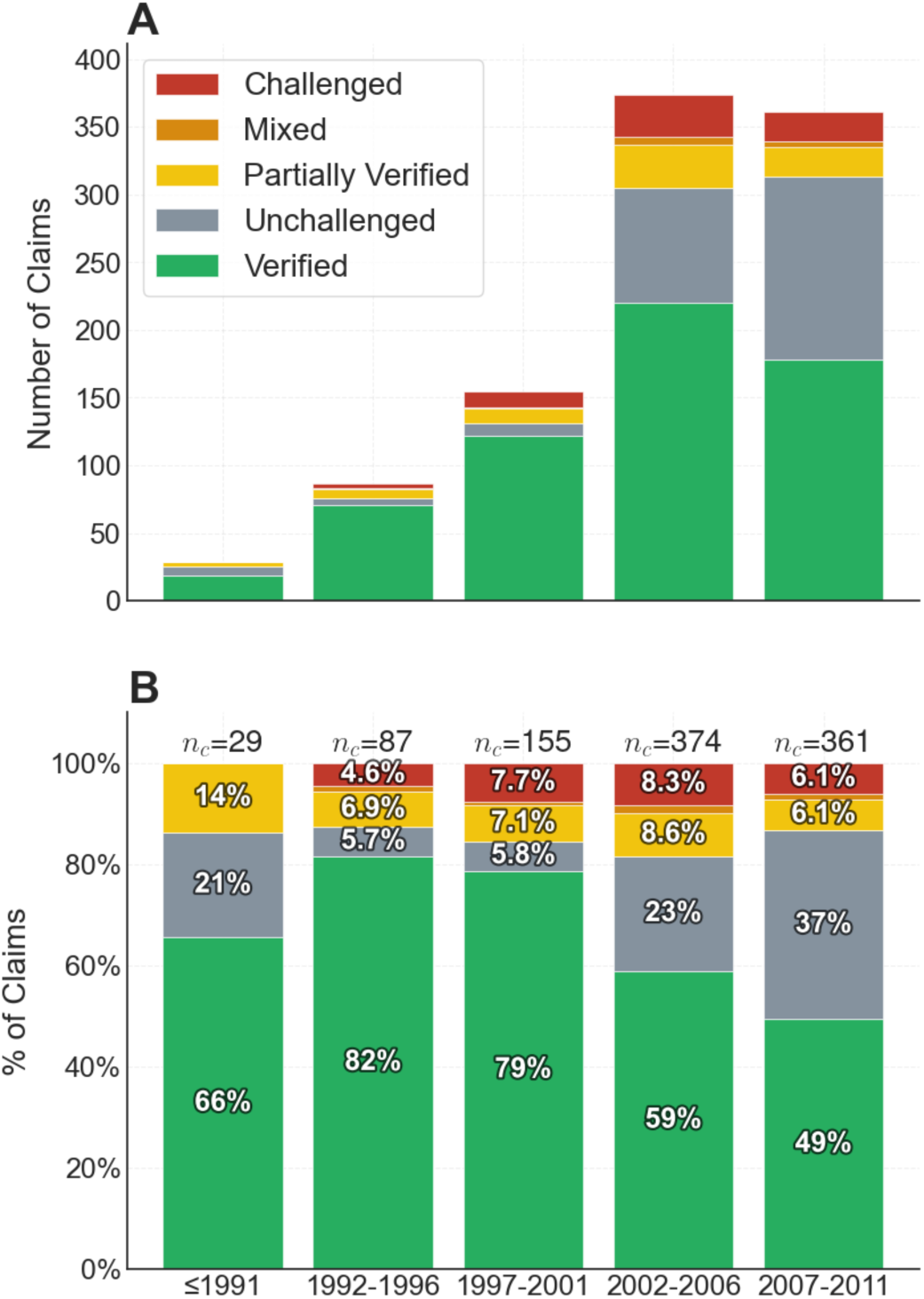
Evolution of claim replicability over time. (A, B) The distribution of claims by categories of assessment over time is shown in A, and the proportion normalized over the total number of claims per time period is shown in B.

Interestingly, we observed a peak in verified claims between 1992 and 2001, accompanied by a lower percentage of unchallenged claims (**Figure 3**). This period coincided with the early expansion of the field when many researchers worked on similar research questions with a relatively narrow conceptual framework of innate immunity. The subsequent increase in unchallenged claims may reflect the rapid conceptual expansion of the field, which likely outpaced the growth in the number of researchers, reducing opportunities for overlapping or validating research efforts. An alternative explanation for this trend is that in the early stages of the field, researchers uncovered the largest, easily detectable effects. As the field matured, focus shifted to subtler phenotypes and complex regulatory mechanisms, making replication more challenging.

Overall, our time-course analysis suggests that irreplicability has increased over the decades, aligning with the reproducibility crisis narrative. However, an alternative hypothesis is that irreplicability varied alongside the field’s popularity, peaking once when the field became popular and had attracted newcomers who may not have been fully acquainted with its established standards or who opportunistically joined the field. Furthermore, the rise in the percentage of challenged claims was accompanied by an increase in unchallenged claims, indicating a reduced degree of validation within the field.

### First-author patterns of irreplicability

Having analyzed the pattern of irreplicability across a large number of articles, representing contributions from multiple laboratories over five decades, we next investigated the distribution of irreplicability among authors and laboratories. A central question is whether irreplicability is evenly distributed across the scientific community, or if hubs — specific authors or laboratories —are responsible for producing more invalid claims. First, we explored whether replicability is associated with the first author, who typically conducts the work, or with the last author, who manages the laboratory and may foster a research environment that produces irreplicable results.

To explore the relationship between irreplicability and first authorship, we computed the percentage of “challenged” articles attributed to each first author. Additional information on first author was obtained through interviews with principal investigators (PIs) or via public LinkedIn profiles, including variables such as gender and position in the laboratory.

Among the 400 articles analyzed by the *ReproSci* project, additional information for a first author was identified for 394 articles, corresponding to 289 unique first authors and 991 major claims. These authors contributed an average of 3.43 claims each. Notably, 46 authors (15.9%, 95% CI: 12.2–20.6%) published at least one non-replicable claim. **Figure 4A** displays the distribution of challenged claims among first authors. The distribution of challenged claims among first authors was highly unequal with a Gini coefficient of 0.881 (inequality index ranging from 0 -perfect equality-to 1 -extreme inequality-, **Figure 4B)**: the top 10% of first authors accounted for 73.9% of challenged claims, and the top 20% accounted for all challenged claims. This inequality exceeded that expected by chance (reassigning randomly the 69 challenged labels across individual claims, while preserving each author’s number of claims, p = 0.000001). It also remained greater than expected when claims from the same paper were kept together (p = 0.015; Supplementary **S9**). The concentration of challenged claims among first authors cannot be explained solely by differences in the number of claims or by clustering within papers. We next investigated whether irreplicability varied by the gender of the first author. The proportions of challenged articles were similar for male and female first authors: 7.3% and 6.7%, respectively, and this difference was not statistically significant (OR = 1.09, 95% CI: 0.66–1.78, Fisher’s exact p = 0.802; **Figure 5A**).

**Figure 4:**
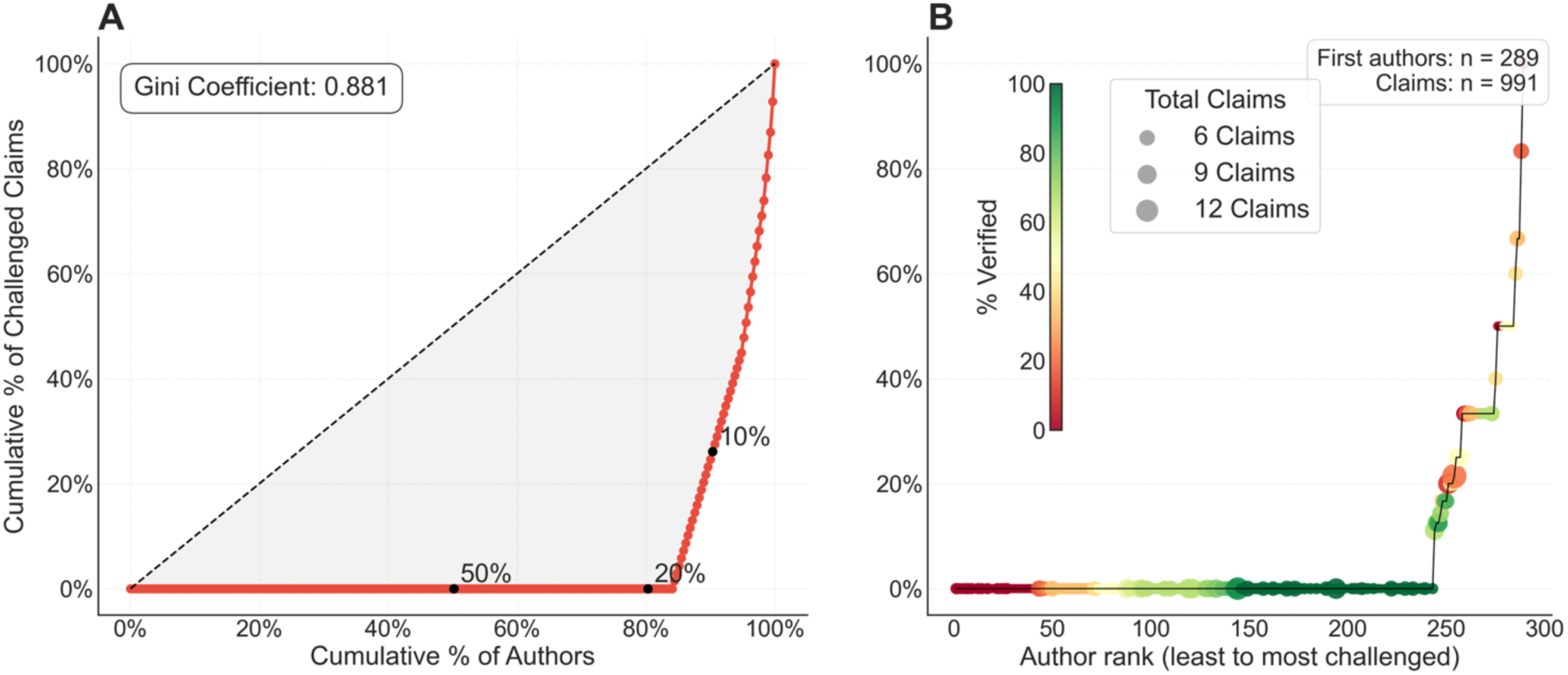
distribution of irreplicability according to first authors. A) Distribution of irreplicable claims among first author. We note that all irreplicable claims are produced by less than 20% of the first authors. Both panel includes 289 first authors contributing 991 claims from 394 articles. Six articles containing 15 claims were excluded because the first author was not identified. B) Distribution of irreplicability for all first authors. Each dot represents an author, ranked according to the proportion of challenged claims. The size of the circle indicates the number of claims and the color the percentage of verified claims. Authors tied on challenged claim proportion are ordered by verified claim proportion.

**Figure 5.**
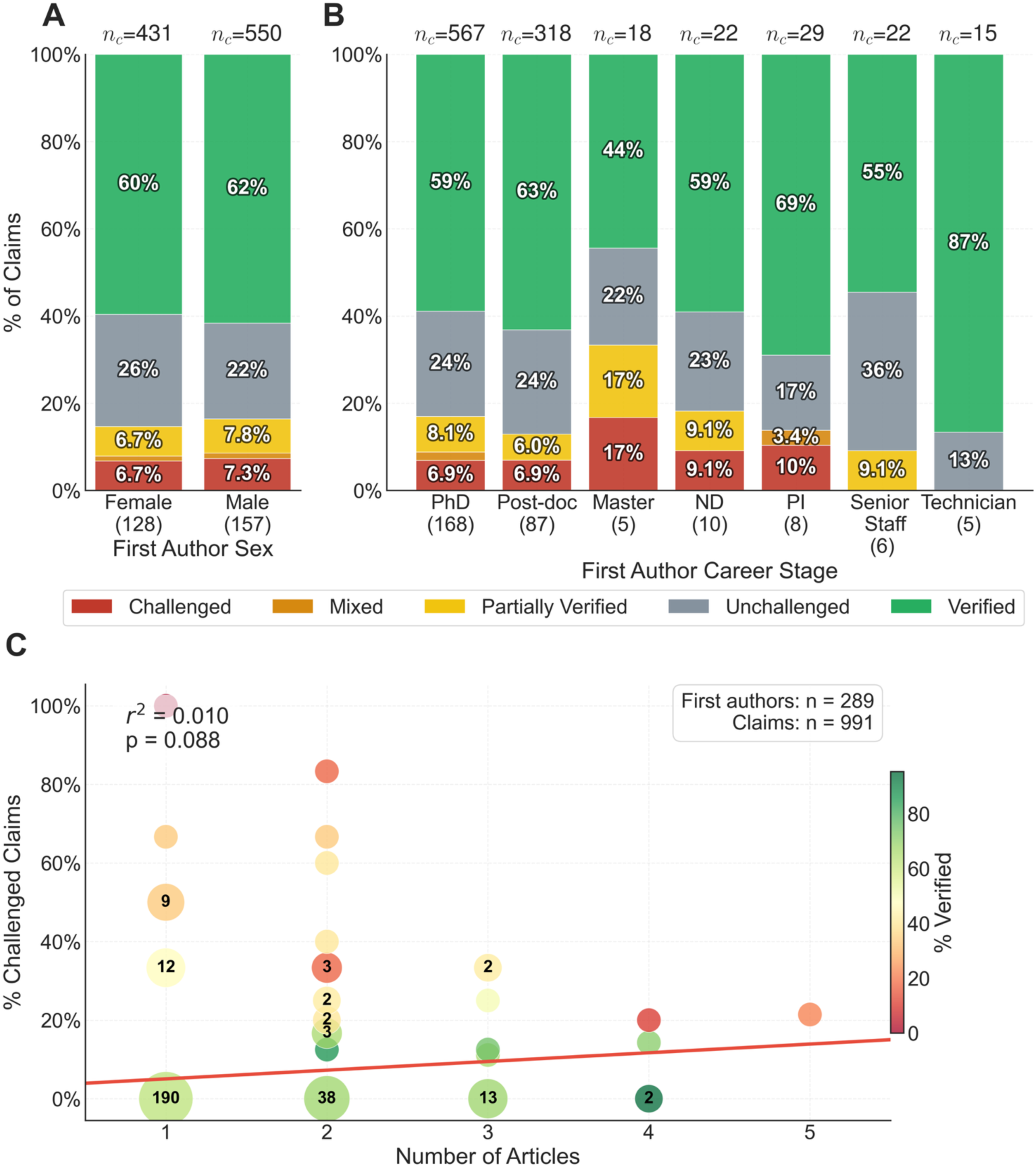
Influence of parameters associated with first author on replicability. A) Repartition of claims in each assessment category according to the gender of the first authors. The gender of four authors, contributing 10 claims, could not be inferred based on first name ambiguity explaining why the total number of authors is 285. B) Repartition of claims in each assessment category according to status of the first authors. The stages of career of 279 first authors were obtained by interviewing last authors, querying Linkedln public profiles or extrapolated according to the publication list. Senior staff refers to permanent position scientist working in a lab. ND stands for Not Determined. C) Proportion of challenged claims as a function of the number of articles published by each of the 289 identified first authors. Coincident observations are combined into a circle. Circle area and the number shown indicate the number of first authors represented. Color indicates the percentage of claims classified as verified.

We were able to determine the career stage of 279 of the 289 identified first authors at the time of the article was published, most being PhD students (n=168) or postdoctoral researchers (n=87), with smaller numbers identified as technicians (n=5), Masters students (n=5), PIs (n=8), or permanent staff (n=6) (**Figure 5B**). No major differences in irreplicability were observed between PhD students and post-docs. While we did observe a higher proportion of challenged claims among PIs and Master’s students, and a lower proportion among technicians and permanent staff, these trends are difficult to interpret due to small sample sizes.

Finally, we tested the relation between the number of articles published by a first author and the proportion of irreplicable research. No significant trend was observed (linear regression; r^2^=0.01, p=0.088; **Figure 5C**), suggesting the level of author productivity is independent of irreplicability. In summary, we observed considerable variability in irreplicability among first authors, suggesting that individual authors may substantially contribute to reproducibility issues.

### Lead authors pattern of irreplicability

We then turned to the question of whether irreplicability varies according to laboratory. As it is common practice in biology, we assigned the last author (referred to as PI for principal investigator) to represent the laboratory. The 400 analyzed articles were co-authored by 156 leading authors from various institutions, predominantly from France, Sweden, Japan, South Korea, and the USA. For each last author, we estimated the number of major claims in each of the five categories. The variation in irreplicability across these PIs is shown in **Figure 6A**. A statistical dispersion analysis in **Figure 6B** revealed that challenged claims were also unevenly distributed among leading authors (Gini coefficient = 0.856): the top 10% accounted for 71.0% of challenged claims and the top 20% accounted for 94.2%. The observed inequality exceeded that expected when challenged labels were reassigned randomly across individual claims (p = 0.00081), but not when claims from the same paper were kept together (p = 0.127; Supplementary **S9**). The apparent concentration among leading authors could be explained by multiple challenged claims arising from the same papers. We next explored whether there were differences in irreplicability between male and female last authors. The percentage of challenged claims was slightly higher for male last authors (7.0%, 95% CI: 5.5%– 9.0%) compared to female last authors (5.8%, 95% CI: 3.1%–10.7%), but this difference was not statistically significant (OR = 1.22, 95% CI: 0.59–2.51; Fisher’s exact p = 0.73; **Figure 7A**).

**Figure 6:**
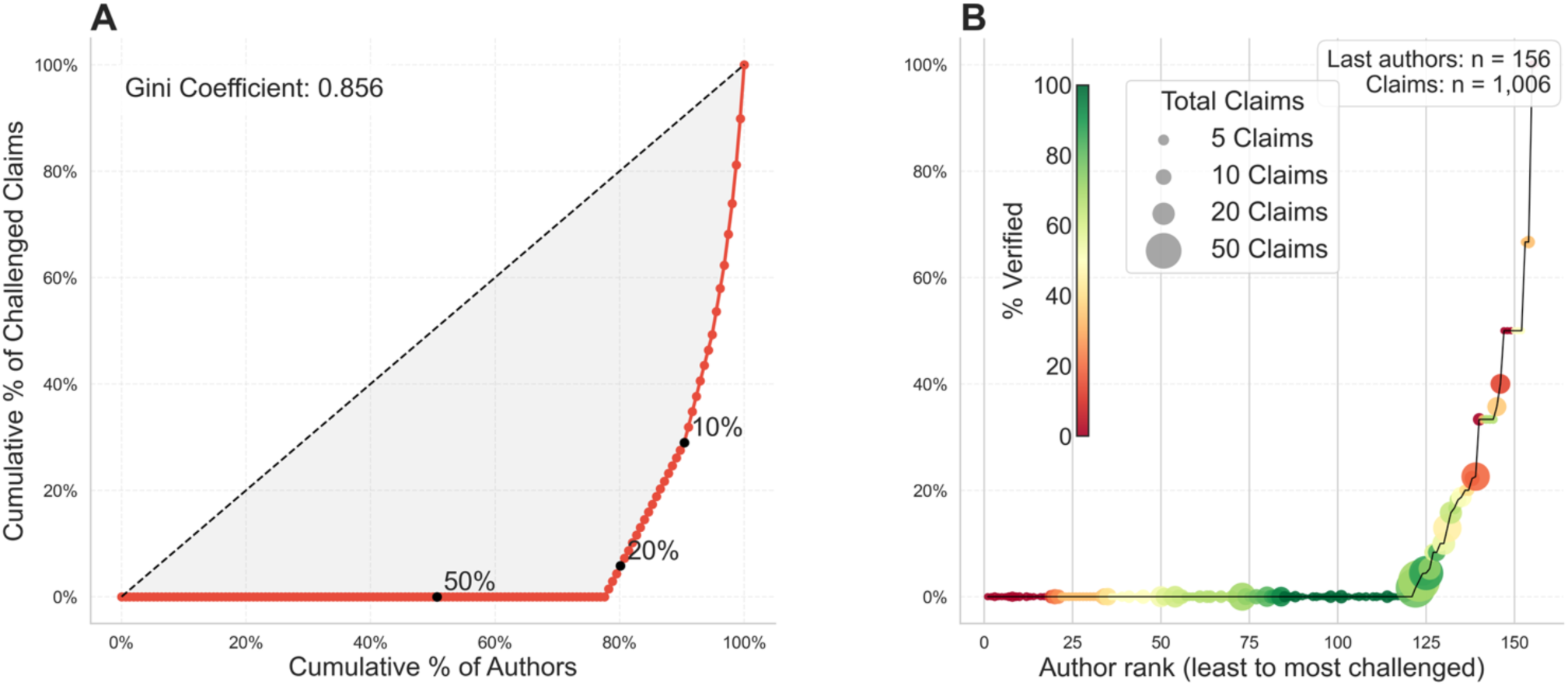
Distribution of irreplicability according to last authors (PI). A) Distribution of irreplicable claims among leading authors. Both panels include all 156 last authors, representing 1,006 claims from 400 articles. B) Distribution of irreplicability for all leading authors. Each dot represents a leading author, ranked according to the proportion of challenged claims. The size of the circle indicates the number of claims and the color the percentage of verified claims. Authors tied on challenged claim proportion are ordered by verified claim proportion.

**Figure 7.**
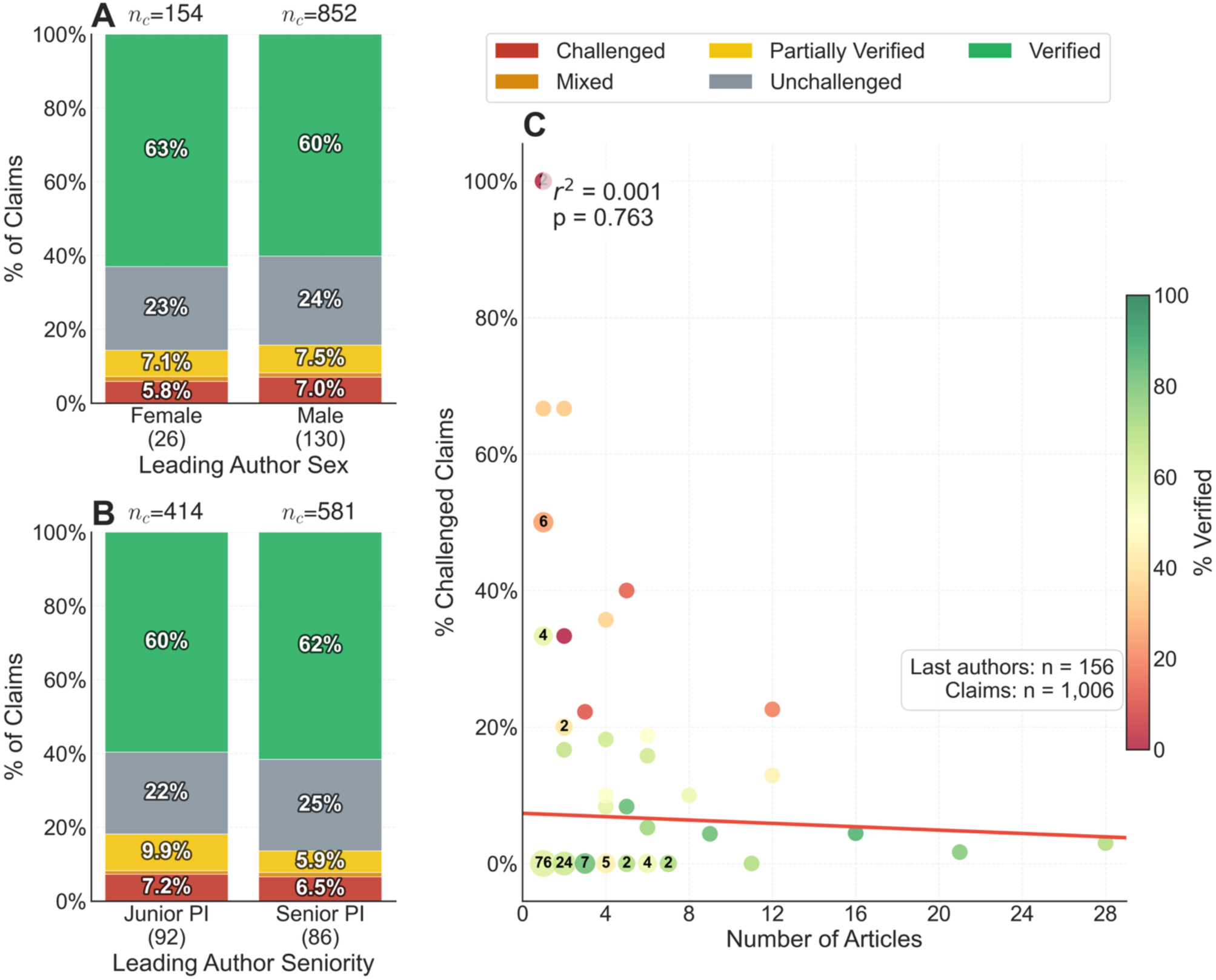
Influence of various parameters associated with the last author on replicability. A) Distribution of claims by category according to the gender of all 156 last authors. B) Distribution of claims by category according to the seniority of the last authors. Senior PI were defined as having published a last author article at least 5 years before the considered articles. The total of junior and senior authors is higher than the total number of last authors as 23 PIs published both as junior and senior. 11 claims from four articles are excluded because they lacked a junior/senior classification. C) Proportion of challenged claims as a function of article count for all 156 last authors. Authors with identical coordinates are combined into a single circle; circle size and internal labels indicate the number of leading authors represented.

Pressure to publish is typically stronger among junior PIs who often lack tenure and must establish their laboratory. To examine whether this pressure might be associated with irreplicability, we classified PIs as junior or senior based on whether they had published a last-author article at least five years prior to the considered publication. Although **Figure 7B** indicates a slightly higher proportion of challenged claims among junior PIs (7.2%, 95% CI: 5.1%–10.2%) compared to senior PIs (6.5%, 95% CI: 4.8%–8.8%), this difference was not statistically significant (OR = 1.12, 95% CI: 0.68–1.83, Fisher’s exact p = 0.70). This modest difference was further attenuated when considering mixed and partially verified claims.

We then explored whether the percentage of irreplicability tended to decline according to the number of published articles as last author (**Figure 7C**). We could hypothesize that PIs that have transiently work on *Drosophila* immunity would have more irreplicable claims, or on the contrary that the most productive laboratories compromise speed over rigor. However, no significant trend was observed between the percentage of challenged claims and the number of articles published by an author (linear regression; r^2^=0.001, p=0.763).

We found that the percentage of irreplicability has increased over time as the field became more popular. We therefore explored if this increase in irreplicability affects all PIs or only new PIs that joined the field at a later time. We show in **Figure 8A** the proportion of challenged claims by last authors according to the publication date of their first article on *Drosophila* immunity, either as last or as first authors. We observed a significant association between irreplicability and the period when principal investigators (PIs) entered the field. Specifically, PIs, who entered the field on 1995 or later, had a significantly higher percentage of challenged claims (8.1%, 95% CI: 6.4%–10.2%) compared to those who entered the field before 1995 (2.6%, 95% CI: 1.2%–5.6%; OR = 3.24, 95% CI: 1.38–7.59, Fisher’s exact p < 0.003).

**Figure 8:**
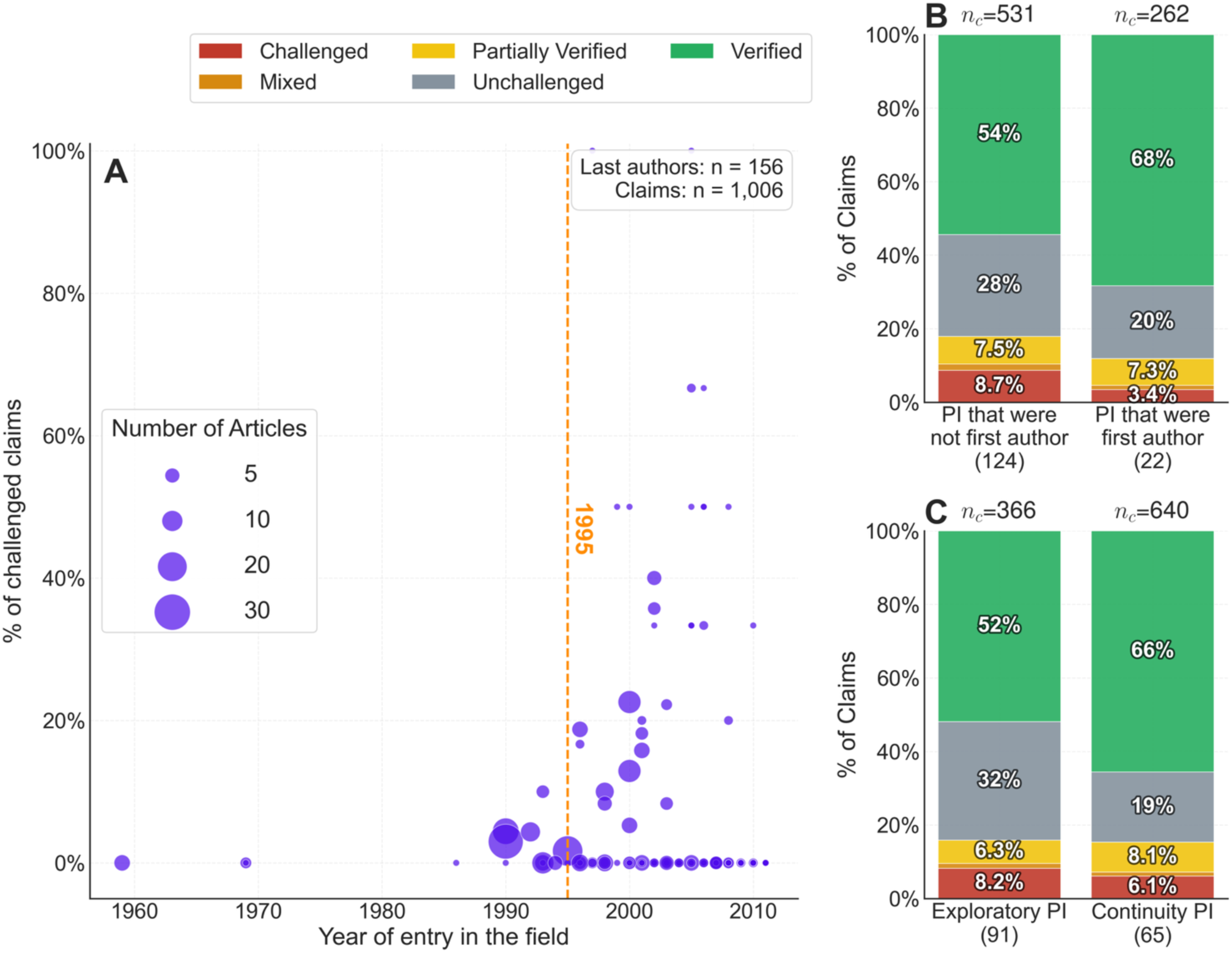
Patterns of irreplicability by last author according to year of entry in the field and research style. A. The proportion of challenged claims by last author is shown according to the time at which the author published their first paper as a first or last authors in the field. It includes all 156 PIs and 1,006 claims; 13 PIs entered the field before 1995 and contributed 227 claims, including 6 challenged claims; 143 entered in 1995 or later and contributed 779 claims, including 63 challenged claims. The increase in irreplicability is associated with PIs whose first publication in the *Drosophila* immunity field, as either a first or last author, occurred in 1995 or later. Circle size indicates the number of published articles by each author. B. Distribution of claims by last author according to whether they have published at least one first author article on *Drosophila* immunity in another laboratory. Prior-experience status was available for 146 PIs. Ten PIs whose first-author publications occurred after they had already become PIs were excluded from comparison. C. Distribution of claims according to research style for all 156 PIs. Proportion of challenged claims by PIs who have continuously work on the *Drosophila* immunity (‘continuity PIs’) or made a transient incursion into the field (‘exploratory PIs’).

Among the PIs working in the *Drosophila* immunity field, we identified 22 who previously published at least one first-author article in this field during their PhD or postdoctoral training in another laboratory. We hypothesized that these authors, having gained prior hands-on experience on *Drosophila* immunity, would be less prone to publishing irreplicable claims. Consistent with this hypothesis, we observed lower levels of irreplicability among these experienced authors (3.4%, 95% CI: 1.8%–6.4%) compared to authors who had not been trained in laboratories working on *Drosophila* immunity (8.7%, 95% CI: 6.6%–11.4%; OR = 2.67, 95% CI: 1.28–5.54, Fisher’s exact p = 0.007; **Figure 8B**).

We conclude that irreplicability varies significantly across laboratories with a limited number of laboratories accounting for most of the irreplicable claims. As for first authors, author gender is not significantly associated with reproducibility. Our studies point to a significant role of experience: PIs that had previously worked in another *Drosophil*a immunity laboratory to the extent that they published a first author article in that laboratory were less prone to publish irreplicable claims.

### Irreplicability according to research styles

Among the various career progression strategies, one involves deepening expertise and specialization within the same field, while another entails transitioning to different fields or exploring different career paths. This may reflect the personality of the scientist, either opting for a more detailed, focused approach or seeking continual new challenges. Scientists from the past were often spending their life on a specific topic within a field. In the last decades, changing field has become the norm (Zeng et al., 2019). Accordingly, scientists are not funded by institutions to work long-term on a single topic anymore, but rather funding is granted by agencies for specific short-term projects. Project grants define research with the primary goal to do ‘breakthrough discoveries’, not accumulative research, which is sometimes treated as the ‘pedestrian’ or the ‘stamp collector’ way of doing research. This leads scientists to move quickly into new research fields in search for opportunities of funding and recognition, a nuanced hypothesis supported by the recent description of a “pivot penalty” for scientists that change career tracks (Hill et al., 2025). We can define these two strategies as either “continuity PIs,” which spend their career on one core topic, and “exploratory PIs”, which regularly venture into new topics. To further explore this question, we classified all last authors as “continuity” PIs or “exploratory” PIs. Exploratory PIs are defined as those who made a transient incursion in the *Drosophila* immunity field before moving on to another field or leaving academia. In contrast continuity PIs worked continuously in the *Drosophila* immunity field throughout their careers. We observed that continuity PIs had a lower proportion of challenged claims (6.1%, 95% CI: 4.5%–8.2%) compared to exploratory PIs (8.2%, 95% CI: 5.8%–11.5%). However, this difference was not statistically significant (OR = 1.38, 95% CI: 0.84–2.26, Fisher’s exact p = 0.24; **Figure 8C**). We also noted a markedly higher share of unchallenged claims among exploratory PIs than among continuity PIs, 32.2 % (95 % CI 27.7%–37.2%) versus 19.1 % (95 % CI 16.2%–22.3%), respectively. This difference was statistically significant (OR = 2.02, 95 % CI 1.50– 2.71; Fisher’s exact *p* < 0.001), reinforcing the idea that researchers who move on to new topics tend to leave a larger fraction of their earlier findings without follow-up or validation by the community. Since unchallenged claims are more likely to be irreplicable, this suggests that irreplicability is higher in the work of PIs with an exploratory style, as opposed to those engaged in a continuity-driven research style.

### Multivariable analysis of predictors of claim irreplicability

We have presented above a comprehensive descriptive data analysis, examining one variable at a time. However, drawing robust conclusions necessitates the joint analysis of all covariates, while appropriately accounting for the non-independence of claims made by the same authors. Thus, we fitted a multivariate Bayesian mixed-effects logistic model to uncover the effect of each covariate, while taking into account all other confounding variables at our disposal (Abril-Pla et al., 2023; Capretto et al., 2022)(**Figure 9**). The model included 869 claims from 345 articles, contributed by 251 first authors and 140 last authors; the remaining 137 claims, including 9 of the 69 challenged claims, were excluded because information was unavailable for at least one variable used in the model.The model simultaneously adjusts for author demographics, laboratory attributes and journal or institutional prestige and predict a binary outcome for each claim: challenged or not challenged (including Verified, Partially Verified, Mixed and Unchallenged). This outcome choice allows us to estimate the probability that a claim has been shown to be irreplicable to date, which combines the probability that it was assessed and the probability that it failed when assessed. As the claims from the same first or last author are not independent, we included a mixed effect for each first author (251) and each last author (140). These effects capture the influence of a particular author in driving a claim reproducibility. We conducted extensive test on model formulation and convergence (see methods, and Supplementary information for posterior predictive and diagnostics plot). As this analysis is Bayesian, we report the Highest Density Interval (HDI) instead of the confidence interval. We observe that none of the first-author factors (gender: OR= 0.94, 94% HDI: 0.41–2.11; trainee stage: OR = 1.16, 94% HDI: 0.49–2.77) or leading-author factors (gender, seniority, continuity, or having entered the field before 1995) showed a decisive association with irreplicability once every other covariate was taken into account (all HDIs span 1). We still observe a trend toward reduction of challenged claims for lead authors having being trained in *Drosophila* immunity or having entered the field before 1995. **Figure S3** shows that lead author with an exploratory career style tends to have significantly more unchallenged claims (OR 2.22, 94% HDI 1.16–4.55). In contrast, university ranking and journal level covariates dominated the model: articles from top-50 universities were almost four times more likely to contain a challenged claim than those from un-ranked institutions (OR 3.68, 1.50–9.28), and papers in trophy journals (Science, Nature, Cell) carried an elevated uncertain risk (OR 1.75, 0.59–4.91) compared to low-impact journals. Intermediate effects for high-impact journals and mid-ranked universities were positive but imprecise. Publication year was modelled as a continuous variable using a three-knot B-spline to allow for non-linear time trends. None of the individual spline coefficients differed credibly from zero, indicating that once journal tier and the other covariates were included the residual calendar-year effect on irreplicability was negligible, indicating that time of publication itself is not a major factor.

**Figure 9.**
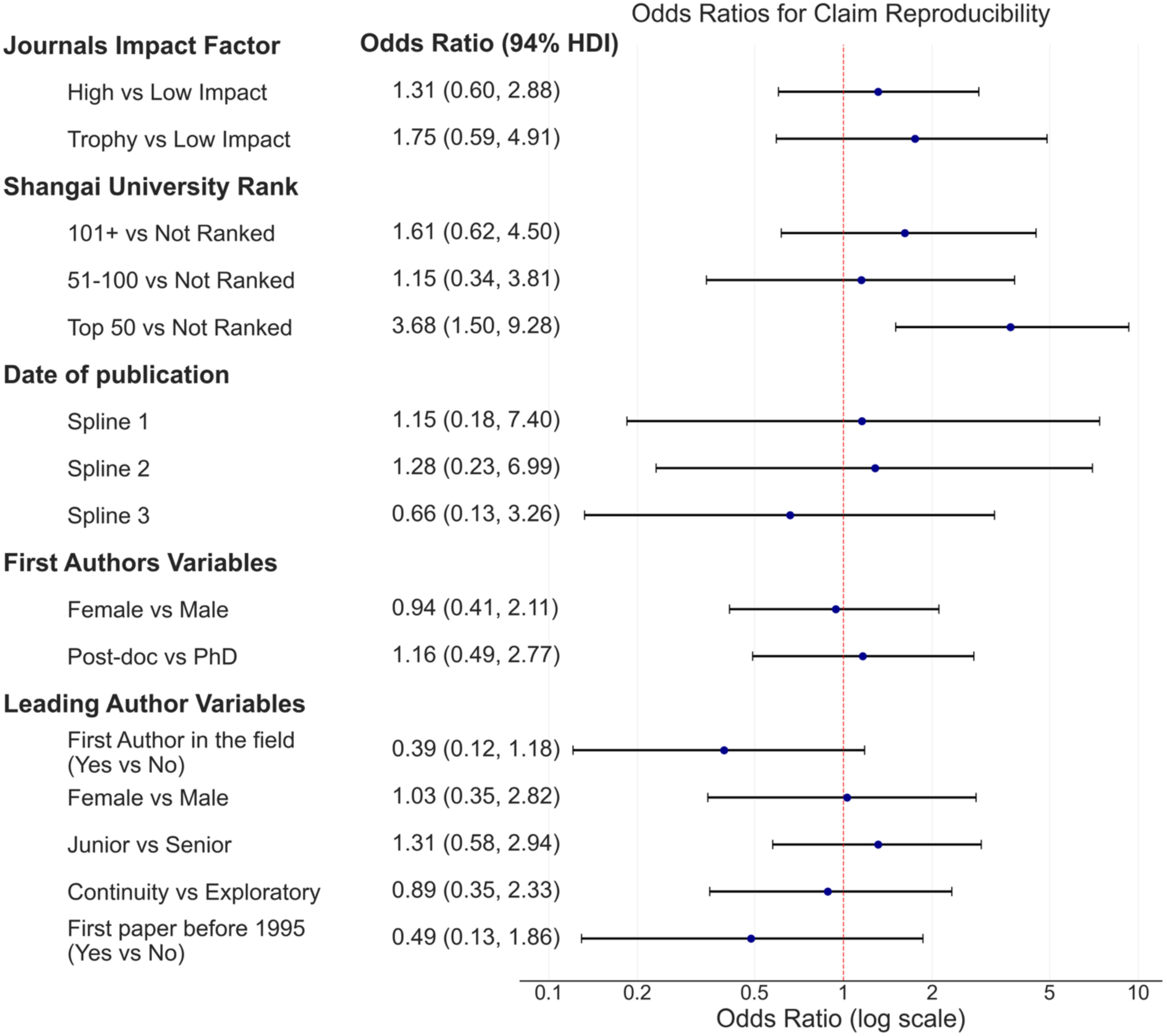
Multivariable analysis of predictors of claim irreplicability. The forest plot reports the odds ratio of each covariate effect on a claim becoming challenged. The multivariate analysis included 869 claims from 345 articles, including 60 challenged claims, contributed by 251 first authors and 140 last authors. Points report adjusted odds ratios and horizontal lines show 94% highest-density intervals.

The hierarchical component comprised 251 random intercepts for first authors and 140 for PIs. The posterior distribution of these intercepts (see Supplementary Information) is broad, confirming that some individuals or laboratories have a systematically higher or lower propensity for challenged claims. Nevertheless, only the three most extreme first-author effects had 95 % highest-density intervals that excluded zero (and all with positive irreplicability) meaning that statistically robust clustering of irreplicability is confined to a handful of contributors; for the vast majority of authors the collected evidence does not support significant differences in replicability from what is expected from a random distribution.

Taken together, the model confirms our descriptive data analysis, namely that it is the prestige of institution, and possibly publication in trophy journals, rather than the personal characteristics of the scientists that best predicts which claims will later prove to be irreplicable.

Similar models to predict unchallenged or not unchallenged claims are described in **Supplementary S3.** Sensitivity analysis excluding unchallenged claims, encoding the impact factor continuously, introducing an article-level effect and using claims status before the validation by the *ReproSci* project are shown in **Supplementary S4**, **S5, S6** and **S7**. Notably, as whether a claim is followed up is not random, we refitted the model excluding unchallenged claims (**Supplementary S4**). This model answers a different question: the probability of failure once a claim has been tested, which removes the detection component but conditions on a selection that depends on the same predictors. The Top-50 universities association was nearly unchanged (OR 3.34, 94% HDI 1.28–9.01), whereas the trophy-journal association was smaller (OR 1.47 vs 1.75). The association with Top-50 universities remains robust in all our analyses except when using only the classifications available before our experimental validation (**Supplementary S7**), although with only 28 challenged claims this analysis has little power.

## Discussion

The *ReproSci* project presents an in-depth expert analysis of reproducibility in an experimental science field with a retrospective period of 14 years. This initiative focuses specifically on *conceptual replicability* —the ability to independently reproduce key findings based on the information provided— rather than direct reproducibility using the exact same data and conditions (Plesser, 2018). Replicability is of critical importance to the scientific community who seeks to build upon published claims. One of our project’s key strengths lies in its comprehensive scope: it spans over fifty years of research, from 1959 to 2011, within a specific scientific community, resulting in a robust database that synthesizes the main findings across decades. This time frame encompasses the formative years of the field, characterized by a small number of pioneering laboratories, followed by a period of rapid expansion driven by increased interest, heightened competition, and publication in high-impact journals. This growth phase was marked by a proliferation of knowledge and conceptual innovation, with novel approaches to studying the immune system emerging. With the awarding of the Nobel Prize in 2011 and subsequently the shift toward more translational research, the growth pace of the *Drosophila* immunity field has since stabilized, and the field remains highly active. Although our study centers on a specific experimental domain at a particular moment in time with a specific geographical development, it reveals broader patterns of irreplicability that are likely generalizable. These findings contribute meaningfully to the ongoing conversation about the reproducibility crisis in the experimental life sciences (Fanelli, 2018; França and Monserrat, 2018; Hunter, 2017; Peng, 2015).

Contrary to the more dramatic narratives surrounding the reproducibility crisis The Brazilian Reproducibility Initiative, 2026; Baker, 2015; Errington et al., 2021), our study shows that a substantial proportion of claims —61%— can be considered as verified. The proportion of claims that have been directly challenged remains relatively low (7%), though likely underestimated. This indicates that research in the *Drosophila* immunity field is rather solid, possibly due to the power of *Drosophila* genetics, the collective ethos of sharing within the community, a less competitive environment, and a lack of influence associated with translational research. Since our study is largely based on published literature, we cannot exclude the possibility that the high replicability rate we report is inflated by confirmation bias in the *Drosophila* immunity field. This bias could arise either because attempts to replicate previous findings tend to converge on similar results due to researcher expectations, or because studies confirming earlier results are more likely to be published (Nissen et al., 2016; Song et al., 2013). However, we consider this explanation unlikely, as the validity of published claims was not accepted uncritically but subjected to informed expert assessment who did not take claims at face value when assessing whether they were validated or challenged. Notably, we find that even highly cited articles—often perceived as particularly robust—contain claims that have later been challenged. Since the website was launched more than two years ago, we have received only three community comments—one on a major claim and two on minor points—prompting us to refine our conclusions. Although this could reflect limited community engagement, it may equally suggest that our assessments are largely accepted. Finally, it should be underlined that the categories we are using in this project do not capture all issues related to replicability, but the conceptual validation of the claims. This difference might explain the higher level of replicability we observed. Overall, our findings are consistent with the view that knowledge of *Drosophila* innate immunity has expanded substantially over recent decades, while also suggesting that scientific findings in this field remain broadly robust.

A novel and important contribution of our work is the identification of a large fraction of *unchallenged* claims —findings that have never been independently confirmed or refuted in the published literature. Current reproducibility projects tend to overlook unchallenged claims. We argue that these claims warrant greater attention, as they are needed to fully understand how science contributes to knowledge.We observed a strong effect of publication date on the proportion of unchallenged claims (**Figure S3**). We hypothesize the low proportion of unchallenged claims around 1992-2001 coincided with the early expansion of the field when many researchers worked on similar research questions. The subsequent increase in unchallenged claims could result from the rapid expansion of the field, limiting the possibility of overlapping research that would validate them. However, the existence of unchallenged claims could also be explained if they are not replicable. For instance, the absence of follow-up studies could reflect authors that are active in the field being aware of the fragility of specific unchallenged claims, and so not pursuing them. The existence of unchallenged claims could also stem from the reluctance of other researchers to publish contradictory statement due to the stigma of publishing negative results, the so-called “file drawer” effect (Rosenthal, 1979). Indeed, many scientists report abandoning verification efforts when unable to reproduce a colleague’s findings, fearing personal conflict or professional retaliation (DeCoursey, 2006; Kannan and Gowri, 2014; Nissen et al., 2016). Our experimental validation demonstrates that a significant proportion of unchallenged claims are, in fact, non-replicable. However, we did not randomly select the unchallenged claims that were to be experimentally tested, and we favored suspicious claims and claims that were easy to experimentally test. Thus, the true rate of reproducibility of unchallenged claims should not be directly extrapolated from our experimental validation attempts. However, we found that *none* of unchallenged, suspicious claims could be replicated. While confirmation bias driven by our initial suspicion cannot be entirely excluded, the absence of follow-up in the published literature for over a decade later supports our interpretation that these findings were not robust. Of note, we considered many of these unchallenged claims were considered suspicious based on informal exchanges within research communities, that reported difficulties in replicating the findings. Most of these non-replicable suspicious claims were published in high-impact or trophy journals (26/31) with great visibility, perhaps contributing to this privately-shared stigma within the community. Taken together, our findings support the notion that “big stories” published without subsequent follow-up may be indicative of irreplicability.

Another takeaway from our experimental validation is that approximately 50% of the unchallenged claims were not previously flagged as problematic failed the experimental validation test regardless. While a claim could remain unchallenged for years because it is valid and no one has anything novel to add to it, our results suggest the status of a claim being “unchallenged”is not a reliable proxy for its validity. If we reclassify the dataset by counting logically consistent unchallenged claims as “verified” and logically inconsistent ones as challenged, and conservatively assume that 50% of the remaining unchallenged claims fall into the latter, we estimate that the true proportion of challenged claims may approach 14%. When we include mixed and partially verified claims, a rough upper-bound estimate suggests that at least 80% of all published claims on *Drosophila* immunity between 1959-2011 are verified or likely to be true leaving 20% of claims either challenged or difficult to interpret. Importantly, challenged claims can include work that was done well, but where the field has advanced in its understanding and can better interpret the original findings.We believe the percentage of challenged claims likely varies across scientific fields, depending on intrinsic challenges such as the complexity of biological processes, methodological variability, and the field’s attractiveness, which may increase competitive pressure. It is also important to emphasize that our study, given this specific categorization, does not account for articles that—despite containing validated claims—could still be considered problematic due to exaggeration or to the use of fragile methodologies.

In a recent editorial, Fiala and Diamendis suggested that reproducibility efforts should focus on high-impact papers as these articles are most deleterious to the scientific community (Fiala and Diamandis, 2018). In our database, we indeed found that “trophy journals”, considered to contain the most impactful science, are more likely to publish articles with challenged claims than high or low-impact, although the difference was not significant. Three intertwined factors can explain why high-impact factor journals have higher rate of non-reproducibility. First, the uncertainty that comes with exploring the frontiers of knowledge may lead to investigations that are founded on an incomplete understanding of the major factors influencing the results. Second, the unconventional nature of some high-impact discoveries, and the fact that claims reported in these journals generally attract more attention, increases the likelihood that other researchers will attempt to replicate them — thereby raising the number of challenged claims. And third, systemic biases that pressure scientists to publish in these journals for prestige might encourage scientists to take shortcuts in finalizing their studies, increasing the chance of challenged claims. Our study shows that claims published in high-impact and trophy journals are less likely to remain unchallenged, which may reflect a greater number of replication attempts on these claims. However, this higher rate of follow-up work cannot fully explain by itself the higher proportion of challenged claims in trophy journals. Compared to low rank and trophy journals, high-ranking journals have the best ratio of verified to challenged claims, as low-impact journals have a high percentage of unchallenged claims.

Strikingly, we observed higher rate of non-replicable claims for scientists working in universities in the top 50 Shanghai ranking. Previous work has suggested journal rank as a major associate of irreplicability (Brembs et al., 2013), and that high-impact journals favour authors from leading institutions (Kulal et al., 2025). Our study highlights how university ranking is among the most striking correlates of irreplicability. Taken together, the cause and effect relationship at play here may be driven most by factors associated with institutional prestige, such as trust among networks of colleagues that act as editors and reviewers, leading to greater acceptance rates at high-impact journals. Importantly, our results suggest this greater acceptance does not strictly reflect a high quality of the work being published, but rather a willingness to trust irreplicable work more frequently if it comes from a prestigious institute. Our study spans a period during which most research was done in North America, Europe, Japan and South Korea. It does not capture the rise of science in China, or rapid growth of scientific output across the global north and south, where scientists in some countries face strong incentive for ‘high impact’ publications (Delgado-López-Cózar et al., 2021; Quan et al., 2017). The fact that the *Drosophila* immunity field monitored in this study developed initially in Sweden and France, which have unique research community characteristics, might affect some our conclusions or limit them to a specific period. Future studies in other areas of research might help to confirm if working in top ranked universities correlates with higher rate of non-replicable science.

Our data align with the broader narrative of increasing rates of non-replicable science. One possible explanation is a general decline in the standards of rigor in recent years in the life sciences. However, a non-mutually exclusive hypothesis is that the rise in irreplicability may reflect the increasing popularity of certain fields, creating greater variance by attracting more researchers, who themselves may not be fully acquainted with the field’s established norms and methodological rigor, or who move opportunistically from field to field to publish striking papers. When research on the gut microbiota gained momentum in the 1990s, it was rapidly positioned as a promising avenue to explain a wide array of diseases. Substantial funding and media attention enabled the swift development of the field. However, many early studies lacked methodological rigor and could not be reproduced (Wesolowska-Andersen et al., 2014). The lower scientific standards tolerated in this emerging area —compared to more established fields— facilitated the publication of high-profile articles, encouraging a rapid influx of new research groups. In this context, scientific novelty and perceived importance often took precedence over precision and reproducibility. A similar but more acute pattern was observed during the SARS-CoV-2 pandemic (Ioannidis et al., 2022; Yeo-Teh and Tang, 2021). Alternatively, this reproducibility crisis might be more linked to specific features of our time and the competitiveness of science, that may lead some scientist to distorting science to favor their own success (Lemaitre, 2016).

Importantly, our analysis shows that irreplicability is not evenly distributed across all researchers. While irreplicability can stem from the inherent difficulty or novelty of certain research questions, our findings point to the significant influence of both the first author (who performs the work) and the laboratory environment. Surprisingly, we did not observe a major effect of experience level: irreplicability rates were similar between PhD students and postdoctoral researchers and did not markedly decrease with the number of publications. Similarly, we only observe a slightly but not significant higher irreplicability rate among junior faculty compared to senior professors. This suggests that irreplicability is not inherent to early career stage but rather to certain research practices and environments. That said, we did observe a higher irreplicability rate among PIs that entered the field from a non-*Drosophila* immunity background. In contrast, we observed a trend toward higher replicability when PIs had been first authors in the laboratories where they received their training in *Drosophila* immunity. This finding is consistent with the value of traditional scientific training pathways, in which early-career researchers develop their expertise within established research environments before gaining independence. Conversely, it suggests that researchers entering a field without such prior experience may face greater challenges in producing replicable findings.

Another noteworthy finding is our suggested link between irreplicability and “continuous” or “exploratory” research style. We observed that authors with a more exploratory research style have slightly higher proportions of challenged claims and a significantly higher proportion of unchallenged claims, the latter being more likely to be challenged as shown by our experimental replication. Exploratory researchers have worked only transiently in the field, often leaving it after its peak period of popularity. In recent years funding agencies and universities favored this exploratory research style focusing on breakthroughs, notably by funding projects that may be rapidly discontinued. As a counterpoint of this, our study highlights the value and reliability of laboratories that tend to maintain long-term commitment to a specific field.

An important aspiration in the field of metascience study would be the ability to predict claims or articles that are likely irreplicable. Our retrospective analysis identifies several criteria that correlate with irreplicability, although with different strengths of evidence. These include i) articles being published by authors working in top-ranked institutions, ii) articles being published in trophy journals, iii) the absence of follow-up studies (i.e. being ‘unchallenged’), and iv) articles being published by authors with an exploratory research style. Of the factors included in the multivariable model, only top-50 institutional ranking was credibly associated with irreplicability across most sensitivity analyses. The trophy-journal and research-style associations were weaker and should be considered tentative. In contrast, having published as first author in another laboratory within the field might be associated with higher replicability (OR 0.39, 94% HDI 0.12–1.18), a trend stronger than for journal tier or research style. The absence of follow-up studies regarding claims published in high-impact journals is likely the most useful proxy to question the replicability of these claims.

### Limitations

This work has several limitations. First, the definition of the claim, the validity, the determination of the validation was influenced by the annotating authors (HW and BL) and should be considered as expert assesment. While the *ReproSci* website was opened to the community, many PIs were reluctant or not interested in the project of testing old claims that represent “scientific cold cases”. Second the leading author is stakeholder of the project having published several articles that were included in the analysis, including 44 articles with B.L. as co-author with 4 as first and 21 as last author. Thus, this may not represent the general view of the community. Third, our conclusions rest largely on retrospective data from the literature. A further limitation is the possible confounding among the predictors of irreplicability themselves (journal tier, institutional ranking, etc.). Moreover, as these predictors influence whether a claim is followed up or not, our estimates of irreplicability partly reflect differences in scrutiny. Finally, the findings we report may be specific to a particular field, model organism, and time period, and may be strongly influenced by factors idiosyncratic to their development. Because this analysis is limited to a single field (immunity) and model system (*Drosophila*), it remains unclear to what extent these findings generalize to other fields, organisms, or more recent studies. However, we believe that despite these limitations the systematic and retrospective analysis of a research field provides insights to better apprehend how science progresses despite errors and the impact of irreplicability.

## Methods

### Selected articles, annotation, experimental validation of the *ReproSci* project

We provide here a succinct method of the *ReproSci* reproducibility project (see the companion article (Westlake et al., 2026) and the ReproSci website for more details). In brief, a list of 400 publications published before 2011 was generated using a curated search string on the publicly available PubMed database. This limitation was imposed to allow a time frame in which follow-up studies may be published that may verify, expand on or contradict the content of the publications selected for the study. Selected primary articles were annotated by a single researcher (HW) and reviewed by a second researcher (BL) to ensure consistency and accuracy across the project. An online database, *ReproSci* (https://ReproSci.epfl.ch/), was constructed to organize the annotations and provide an interactive and searchable interface. Claims were extracted through careful reading of the selected primary articles and organized according to their importance within the paper: a single Main claim representing the title claim of the paper or overarching finding; 3-4 Major claims emphasized in the abstract or those contributing to the overarching finding; and a variable number of Minor claims representing other findings of note not emphasized in the abstract. Claims were cross-checked with evidence from previous, contemporary and subsequent publications and assigned a verification category (see **Table 1**). A short statement explaining the evidence and citing sources was written to justify the categorization and provide additional context for the claim; this assessment was linked to each claim and can be viewed on the *ReproSci* website. The category ‘unchallenged logically consistent” and ‘‘unchallenged logically inconsistent’ refer to claims that although not directly verified are corroborated or not in with current literature (see *ReproSci* for justification) When two articles reported the same main claim that was judged as “verified,” each instance was counted in the analysis. However, when validation came from studies published after 2011, the claim was counted only once. Thus, our study specifically evaluates the reproducibility of claims originating from a defined period (pre-2011). A subset of unchallenged claims (45 major and 11 minor) was selected for direct experimental reproduction by various authors (see author list) from nine laboratories with relevant expertise. Unchallenged claims that were considered as ‘suspicious’ are claims that were already tested in the host laboratory without success, claims whose validity are debated at meeting, claims from high-ranked journal with no follow-up. This project is concerned with the accuracy of the conclusions themselves rather than the reliability of methods, so we employed conceptual replicability or indirect or inferential reproducibility, where experimental procedures that may be different from those originally employed are used to independently verify the claim. Thus, an article’s scientific value does not necessarily align with whether it has been verified or challenged, and our categorical framework does not capture all issues related to reproducibility. The *ReproSci* website was made available to the *Drosophila* scientific community on July 11, 2023, with an email to encourage to comment on the annotations and verifications and contribute evidence.

### First authors and last author classification

Information regarding the gender and status of first authors were either obtained by interviewing PIs (for 191 first authors) or found on publicly available websites (LinkedIn and university websites) or were implied based on their first name usage (gender) and their PubMed publication lists (for the other 100 first authors). When some of these information could not be retrieved, these authors were not included in the analysis.

PIs were manually classified as senior or junior whether they have or have not published a last author article in PubMed at least five years before the considered publication. Continuity and exploratory PIs were manually classified according to their PubMed publication list. Continuity PIs are PIs that have continued to work in the *Drosophila* immunity field until now or to the end of their scientific career (deceased, retired, left academia). Exploratory PIs are PIs that have done a transient incursion in the *Drosophila* immunity field, moving to other topics of research.

### Statistical analysis

The analysis was done on the 1006 major claims of the 400 articles. The cleaned database contains one row per major claim with linked article-, author- and laboratory-level covariates, which was subsequently analyzed. Every output, counts, proportions and graphics, exploratory summaries and Bayesian mixed-effects models are defined under a reproducible Python workflow, which enable the full replication of the main results and every figure in the manuscript.

The ninety-five-percent confidence intervals (CI) for single proportions were determined using Wilson’s score method. Whenever a binary predictor (e.g. author gender) was cross-classified with the binary outcome “claim challenged”, we reported the odds ratio with its 95% CI and the two-sided *p*-value from Fisher’s exact test; The raw contingency tables for every binary predictor appear in the Supplementary document.

We compared the observed Gini coefficients with two permutation analyses, each based on 1,000,000 permutations. First, we randomly reassigned the challenged labels across individual claims while keeping every claim attached to its original author. This preserved the number of claims contributed by each author. Second, we kept the challenged-claim pattern of each paper together and reassigned these patterns among papers containing the same number of claims. This additionally accounted for the possibility that claims from the same paper were challenged because of a shared problem. For each permutation, we recalculated the Gini coefficient across authors and calculated the p-value as the proportion of simulated coefficients that equalled or exceeded the observed coefficient. Full results are reported in **Supplementary Document**.

### Multivariable modelling

Two Bayesian regressions were fitted with weakly-informative priors. Posterior summaries are presented as odds ratios (OR) with 94 % highest-density intervals (HDI).

**1.** Hierarchical logistic regression for claim irreplicability The outcome was whether a major claim was later classified as challenged. Fixed-effect predictors captured article prestige, author characteristics and laboratory context: journal tier (Low, High, Trophy), university-ranking tier (Not ranked, 101 +, 51–100, Top 50), publication year modelled with a three-knot B-spline, first-author gender and training stage (PhD vs post-doc), last-author gender, laboratory seniority (junior vs senior PI), continuity style (continuous vs exploratory) and whether the laboratory had published in the field before 1995. Claims from one author not independent, so random intercepts were included for both first-author key and last-author key.

All models were estimated with the dynamic-HMC NUTS sampler (target _accept = 0.9); four independent chains of 4,000 tuning iterations and 4,000 draws were run for every model. Convergence was confirmed both visually and numerically (all Rhat≤1.01; all effective sample sizes > 400). Supplementary Fig. 1 posterior-predictive checks; Exact formulas, priors, sampling statements and notebooks are available in the public repository (**Supplementary Document**).

## Acknowledgments

We thank the BioInformatics Competence Center of EPFL-UNIL for generating the *ReproSci* website. This work was supported by Swiss National Science Foundation 310030_189085 and the ETH-Domain’s Open Research Data (ORD) Program (2022). We thank Florent Masson and Samuel Rommelaere for comments on the manuscript.

## Supplementary Information

A supplementary document provides information on the full code needed to reproduce the analysis (S1); the multivariate model, with its convergence diagnostics and author-level random effects (S2, Figures S1 and S2); the multivariate model using unchallenged as the outcome (S3, Figure S3); four sensitivity analyses of the multivariate model: excluding unchallenged claims (S4, Figure S4), encoding the impact factor continuously (S5, Figure S5), using the article rather than the claim as the unit of analysis (S6, Figures S6 and S7), and using claim classifications as they stood before experimental validation by the *ReproSci* project (S7, Figure S8); tables of the odds ratios and values for the categorical variables shown in the main text, for first and last authors (S8); the permutation analysis of the Gini index (S9); other categorical comparisons (S10); and a table of journals with impact factor and claim assessment (S11, Table S1).

## References

Abril-Pla O, Andreani V, Carroll C, Dong L, Fonnesbeck CJ, Kochurov M, Kumar R, Lao J, Luhmann CC, Martin OA, Osthege M, Vieira R, Wiecki T, Zinkov R. 2023. PyMC: a modern, and comprehensive probabilistic programming framework in Python. PeerJ Computer Science 9:e1516. DOI: 10.7717/peerj-cs.1516

Amaral et al.,. 2025. Estimating the replicability of Brazilian biomedical science. bioRxiv 2025.04.02.645026.

Amaral OB, Neves K, Wasilewska-Sampaio AP, Carneiro CF. 2019. The Brazilian Reproducibility Initiative. eLife 8:e41602. DOI: 10.7554/eLife.41602

Baker M. 2016. 1,500 scientists lift the lid on reproducibility. Nature 533:452–454. DOI: 10.1038/533452a, PMID: 27225100

Baker M. 2015. Over half of psychology studies fail reproducibility test. Nature nature.2015.18248. DOI: 10.1038/nature.2015.18248

Begley CG, Ioannidis JPA. 2015. Reproducibility in Science: Improving the Standard for Basic and Preclinical Research. Circulation Research 116:116–126. DOI: 10.1161/CIRCRESAHA.114.303819

Brembs B, Button K, Munafò M. 2013. Deep impact: unintended consequences of journal rank. Frontiers in Human Neuroscience 7. DOI: 10.3389/fnhum.2013.00291

Capretto T, Piho C, Kumar R, Westfall J, Yarkoni T, Martin OA. 2022. Bambi: A simple interface for fitting Bayesian linear models in Python. DOI: 10.48550/arXiv.2012.10754

Chalmers I, Glasziou P. 2009. Avoidable Waste in the Production and Reporting of Research Evidence. 114:5.

Da Costa GG, Neves K, Amaral OB. 2022. Estimating the replicability of highly cited clinical research (2004-2018) (preprint). Epidemiology. DOI: 10.1101/2022.05.31.22275810

DeCoursey TE. 2006. It’s difficult to publish contradictory findings. Nature 439:784–784. DOI: 10.1038/439784b

Delgado-López-Cózar E, Ràfols I, Abadal E. 2021. Letter: A call for a radical change in research evaluation in Spain. El profesional de la información. DOI: 10.3145/epi.2021.may.09

Devezer B, Buzbas EO. 2023. Rigorous exploration in a model-centric science via epistemic iteration. Journal of Applied Research in Memory and Cognition 12:189–194. DOI: 10.1037/mac0000121

Eisner DA. 2018. Reproducibility of science: Fraud, impact factors and carelessness. Journal of Molecular and Cellular Cardiology 114:364–368. DOI: 10.1016/j.yjmcc.2017.10.009

Errington TM, Denis A, Perfito N, Iorns E, Nosek BA. 2021. Challenges for assessing replicability in preclinical cancer biology. eLife 10:e67995. DOI: 10.7554/eLife.67995

Errington TM, Iorns E, Gunn W, Tan FE, Lomax J, Nosek BA. 2014. An open investigation of the reproducibility of cancer biology research. eLife 3. DOI: 10.7554/eLife.04333

Fanelli D. 2018. Opinion: Is science really facing a reproducibility crisis, and do we need it to? Proceedings of the National Academy of Sciences 115:2628–2631. DOI: 10.1073/pnas.1708272114

Fiala C, Diamandis EP. 2018. Benign and malignant scientific irreproducibility. Clinical Biochemistry 55:1–2. DOI: 10.1016/j.clinbiochem.2018.03.015

Fidler F, Chee YE, Wintle BC, Burgman MA, McCarthy MA, Gordon A. 2017. Metaresearch for Evaluating Reproducibility in Ecology and Evolution. BioScience biw159. DOI: 10.1093/biosci/biw159

Fidler F, Wilcox J. 2018. Reproducibility of scientific results.

Fortunato S, Bergstrom CT, Börner K, Evans JA, Helbing D, Milojević S, Petersen AM, Radicchi F, Sinatra R, Uzzi B, Vespignani A, Waltman L, Wang D, Barabási A-L. 2018. Science of science. Science 359. DOI: 10.1126/science.aao0185

França TF, Monserrat JM. 2018. Reproducibility crisis in science or unrealistic expectations? EMBO reports 19:e46008. DOI: 10.15252/embr.201846008

Goodman SN, Fanelli D, Ioannidis JPA. 2016. What does research reproducibility mean? Science Translational Medicine 8:341ps12–341ps12. DOI: 10.1126/scitranslmed.aaf5027

Hill R, Yin Y, Stein C, Wang X, Wang D, Jones BF. 2025. The pivot penalty in research. Nature 1–8. DOI: 10.1038/s41586-025-09048-1

Hunter P. 2017. The reproducibility “crisis.” EMBO reports 18:1493–1496. DOI: 10.15252/embr.201744876

Ioannidis JPA, Allison DB, Ball CA, Coulibaly I, Cui X, Culhane AC, Falchi M, Furlanello C, Game L, Jurman G, Mangion J, Mehta T, Nitzberg M, Page GP, Petretto E, van Noort V. 2009. Repeatability of published microarray gene expression analyses. Nature Genetics 41:149–155. DOI: 10.1038/ng.295

Ioannidis JPA, Bendavid E, Salholz-Hillel M, Boyack KW, Baas J. 2022. Massive covidization of research citations and the citation elite. Proceedings of the National Academy of Sciences 119:e2204074119. DOI: 10.1073/pnas.2204074119

Kannan S, Gowri S. 2014. Contradicting/negative results in clinical research: Why (do we get these)? Why not (get these published)? Where (to publish)? Perspectives in Clinical Research 5:151–153. DOI: 10.4103/2229-3485.140546, PMID: 25276623

Kulal A, N A, Shareena P, Dinesh S. 2025. Unmasking Favoritism and Bias in Academic Publishing: An Empirical Study on Editorial Practices. Public Integrity 1–22. DOI: 10.1080/10999922.2024.2448875

Lemaitre B. 2016. An Essay on Science and Narcissism : Why do high-ego personalities drive research in life sciences?, EPFL press.

Lemaitre B. 2004. The road to Toll. Nature Reviews Immunology 4:521–527. DOI: 10.1038/nri1390

Lesperance DNA, Broderick NA. 2021. Gut Bacteria Mediate Nutrient Availability in Drosophila Diets. Applied and Environmental Microbiology 87.

Macleod MR, Michie S, Roberts I, Dirnagl U, Chalmers I, Ioannidis JPA, Salman RA-S, Chan A-W, Glasziou P. 2014. Biomedical research: increasing value, reducing waste. The Lancet 383:101–104. DOI: 10.1016/S0140-6736(13)62329-6

Meng X-L. 2020. Reproducibility, Replicability, and Reliability. Harvard Data Science Review 2. DOI: 10.1162/99608f92.dbfce7f9

Nissen SB, Magidson T, Gross K, Bergstrom CT. 2016. Publication bias and the canonization of false facts. eLife 5:e21451. DOI: 10.7554/eLife.21451

Peng R. 2015. The reproducibility crisis in science: A statistical counterattack. Significance 12:30–32. DOI: 10.1111/j.1740-9713.2015.00827.x

Piller C. 2022. Blots on a field? Science (New York, NY) 377:358–363.

Plesser HE. 2018. Reproducibility vs. Replicability: A Brief History of a Confused Terminology. Frontiers in Neuroinformatics 11. DOI: 10.3389/fninf.2017.00076

Prinz F, Schlange T, Asadullah K. 2011. Believe it or not: how much can we rely on published data on potential drug targets? Nature Reviews Drug Discovery 10:712–712. DOI: 10.1038/nrd3439-c1

Quan W, Chen B, Shu F. 2017. Publish or impoverish: An investigation of the monetary reward system of science in China (1999-2016). Aslib Journal of Information Management 69:486–502. DOI: 10.1108/ajim-01-2017-0014

Rosenthal R. 1979. The file drawer problem and tolerance for null results. Psychological Bulletin 86:638–641. DOI: 10.1037/0033-2909.86.3.638

Shiffrin RM, Börner K, Stigler SM. 2018. Scientific progress despite irreproducibility: A seeming paradox. Proceedings of the National Academy of Sciences 115:2632–2639. DOI: 10.1073/pnas.1711786114

Song F, Hooper L, Loke YK. 2013. Publication bias: what is it? How do we measure it? How do we avoid it? Open Access Journal of Clinical Trials 5:71–81. DOI: 10.2147/OAJCT.S34419

Steneck NH. 2011. The dilemma of the honest researcher. EMBO reports 12:745–745. DOI: 10.1038/embor.2011.134

Stern AM, Casadevall A, Steen RG, Fang FC. 2014. Financial costs and personal consequences of research misconduct resulting in retracted publications. eLife 3:e02956. DOI: 10.7554/eLife.02956

Twenge JM, Campbell WK. 2009. The narcissism epidemic: Living in the age of entitlement. Simon and Schuster.

Udesky L. 2025. ‘Publish or perish’ culture blamed for reproducibility crisis. Nature. DOI: 10.1038/d41586-024-04253-w

van Wesel M. 2016. Evaluation by Citation: Trends in Publication Behavior, Evaluation Criteria, and the Strive for High Impact Publications. Science and Engineering Ethics 22:199–225. DOI: 10.1007/s11948-015-9638-0

Wesolowska-Andersen A, Bahl MI, Carvalho V, Kristiansen K, Sicheritz-Pontén T, Gupta R, Licht TR. 2014. Choice of bacterial DNA extraction method from fecal material influences community structure as evaluated by metagenomic analysis. Microbiome 2:19. DOI: 10.1186/2049-2618-2-19

Westlake H, David FPA, Tian Y, Krakovic K, Dolgikh A, Juravlev L, Esmangart de Bournonville T, Carboni A, Melcarne C, Shan T, Wang Y, Mu Y, Kotwal A, Pirko N, Boquete J-P, Schüpfer F, Rommelaere S, Poidevin M, Liu Z, Kondo S, Ratnaparkhi GS, Chakrabarti S, Liu G, Masson F, Xiaoxue, Hanson MA, Haobo J, Di Cara F, Kurant E, Lemaitre B. 2026. Reproducibility of Scientific Claims in Drosophila Immunity: A Retrospective Analysis of 400 Publications. eLife 15:RP108404 10.7554/eLife.108404.1

Westlake H, Hanson MA, Lemaitre B. 2024. The Drosophila Immunity Handbook. EPFL press.

Yeo-Teh NSL, Tang BL. 2021. An alarming retraction rate for scientific publications on Coronavirus Disease 2019 (COVID-19). Accountability in Research 28:47–53. DOI: 10.1080/08989621.2020.1782203, PMID: 32573274

Youyou W, Yang Y, Uzzi B. 2023. A discipline-wide investigation of the replicability of Psychology papers over the past two decades. Proceedings of the National Academy of Sciences 120:e2208863120. DOI: 10.1073/pnas.2208863120

Zeng A, Shen Z, Zhou J, Fan Y, Di Z, Wang Y, Stanley HE, Havlin S. 2019. Increasing trend of scientists to switch between topics. Nature Communications 10:3439. DOI: 10.1038/s41467-019-11401-8

