## Supplement file for "A retrospective analysis of 400 publications reveals patterns of irreplicability across an entire life sciences research field"

**Table of content**


1. Reproduction ... 1
2. Multivariate model ... 2
3. Multivariate model using Unchallenged as outcome ... 5
4. Multivariate model excluding unchallenged claims ... 6
5. Multivariate model with continuous impact factor ... 7
6. Multivariate model with article level claims ... 8
7. Multivariate model using pre-experimental classifications ... 10
8. Categorical comparison tables ... 11
9. Permutation analysis of the Gini index ... 12
10. Other comparisons ... 12
11. Table of journals with 2022 impact factor, and claim assessment ... 15
12. References ... 16


#### 1. Reproduction

The full code to reproduce the study is available on https://github.com/jcblemai/drosophila-reproducibility-VOR. There are two options to reproduce all figures and number in the paper. The simpler would be to start from the data in csv format stored in the preprocessed_data/ in the repository. However, one could run the preprocessing manually by:

- Downloading a SQL dump of the ReproSci database (https://reprosci.epfl.ch) and, running the jupyter notebook preprocess_db.ipynb
- Then, getting all manually registered covariates from the excel files using notebook preprocess_xlsx.ipynb.

These two notebooks recreate the folder processed_data/ as it is, correcting some error and using the best source of information for each covariate. This folder contains:

- article_db.csv: data for each article (journal, year) from the ReproSci database
- author_db.csv : data for each author (sex, …) from the ReproSci database
- claims_db_truncated.csv: the main file, over which the two above are merged, has one line per claim, with every other information about its author and articles, from the reprosci database.

- first_author_claims.csv: all covariates from manual collection from first author, merged with the claim file above.
- last_author_claims.csv: same for leading author

The analysis is contained in 4 scripts, that often call helper module for plotting (plot_info.py), data wrangling (wrangling.py) and statistical analysis (stat_lib.py):

- analysis_claims.py generates the analysis of claim data, Main Text figures 1,2 and 3 along with diagnosis information
- analysis_authors_first.py analysis the first author data and generate main text figures 4 and 5
- analysis_authors_last.py analysis the last author data and generate main text figures 6 and 7
- statistical_analysis.py run the statistical analysis of the multivariate model, used to produce main text figure 9.

Running all these scripts will produce all figures, csv files containing categorical odd-ratio, confidence interval and everything used in the main text.

#### 2. Multivariate model

The model formula is:
*challenged_flag ~*
*C(journal_category, Treatment('Low Impact')) + year_s1 + year_s2 + year_s3 + C(F_and_L, Treatment('False')) + C(ranking_category, Treatment('Not Ranked')) + C(First_Author_Sex, Treatment('Male')) + C(PhD_Postdoc, Treatment('PhD')) + C(Leading_Author_Sex, Treatment('Male')) + C(Junior_Senior, Treatment('Senior PI')) + C(Continuity, Treatment(False)) + C(first_paper_before_1995, Treatment(False)) + (1 | first_author_key) + (1 | leading_author_key)*

Where we model publication year was modeled continuously as a 3 knots spline. *(1 | first_author_key*) and *(1 | leading_author_key)* in the formula add independent random intercepts for each first author and each last author. The multivariable analysis included 869 claims from 345 articles, contributed by 251 first authors and 140 last authors; the remaining 137 claims, including 9 of the 69 challenged claims, were excluded because information was unavailable for at least one variable used in the model.

##### Convergence Diagnosis

The multivariate Bayesian logistic-regression model (main text figure 9) was formulated using Bambi (Capretto et al., 2022) and fitted in PyMC (Abril-Pla et al., 2023) NUTS sampler using four chains and 8 000 posterior draws (4 000 for tuning the Hamiltonian sampler + 4 000 draws retained per chain).

Diagnostic summaries indicate that the sampler converged and the model captures the data well:

- R-hat: All parameters had R̂-hat values between 1.00 and 1.01. Because R̂ -hat measures the ratio of within-chain to between-chain variance, values below 1.05 (ideally < 1.01) show the four MCMC chains mixed thoroughly and reached a common stationary distribution.

- Effective sample size (ESS): ESS ranged from 867 to 12 623 across parameters, far exceeding the heuristic minimum of 400. A larger ESS means that the posterior summaries are being estimated from many effectively independent draws, reducing Monte-Carlo error.

- Posterior predictive checks: The posterior predictive is an important part of the Bayesian workflow (Grinsztajn et al., 2021) and verify that data simulated with the fitted model correspond to the observations. The model predicts a mean challenged-claim rate of 7.1 %, closely matching the observed 6.9 %. This alignment suggests the fitted model reproduces the overall frequency of irreplicable claims and is not over- or under-fitting in aggregate. The distribution of posterior predictive check in figure S1.

As additional verification, have also manually verified that omission of covariates does not change the model conclusion, and computed Leave-one-out Cross validation and pareto-k diagnostics. Taken together these diagnostics confirm that the MCMC sampling is stable and that the model provides an adequate description of the data for inference and comparison.

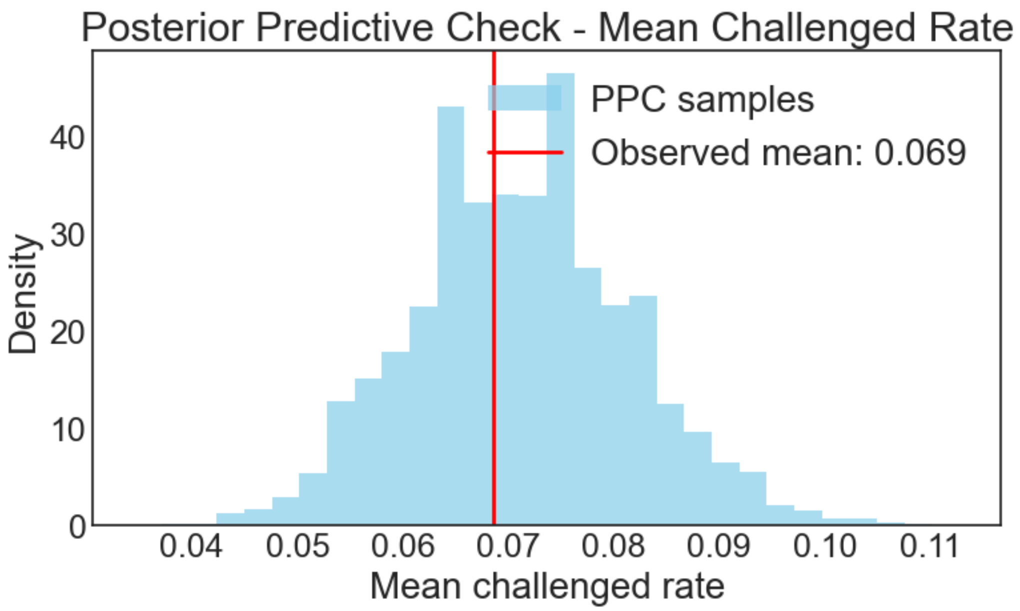


*Figure S1. Distribution of posterior predictive values against the observed mean challenged rate across claims.*

#### Random effects

The terms (1 | first_author_key) and (1 | leading_author_key) in the formula add independent random intercepts for each first author and each last author. These intercepts capture unobserved heterogeneity within authors, acknowledge that claims contributed by the same individual are not statistically independent, and allow the baseline probability of a claim being challenged to vary across authors.  By modelling this clustering explicitly, fixed-effect estimates are protected from inflation of Type-I error and the credible intervals properly reflect both within- and between-author uncertainty. We report in figure S2, for both first and last author random effect, the top and bottom 15 random effect. We see that the only significant effects are increased irreplicability for the top 3 first author with the highest random effects

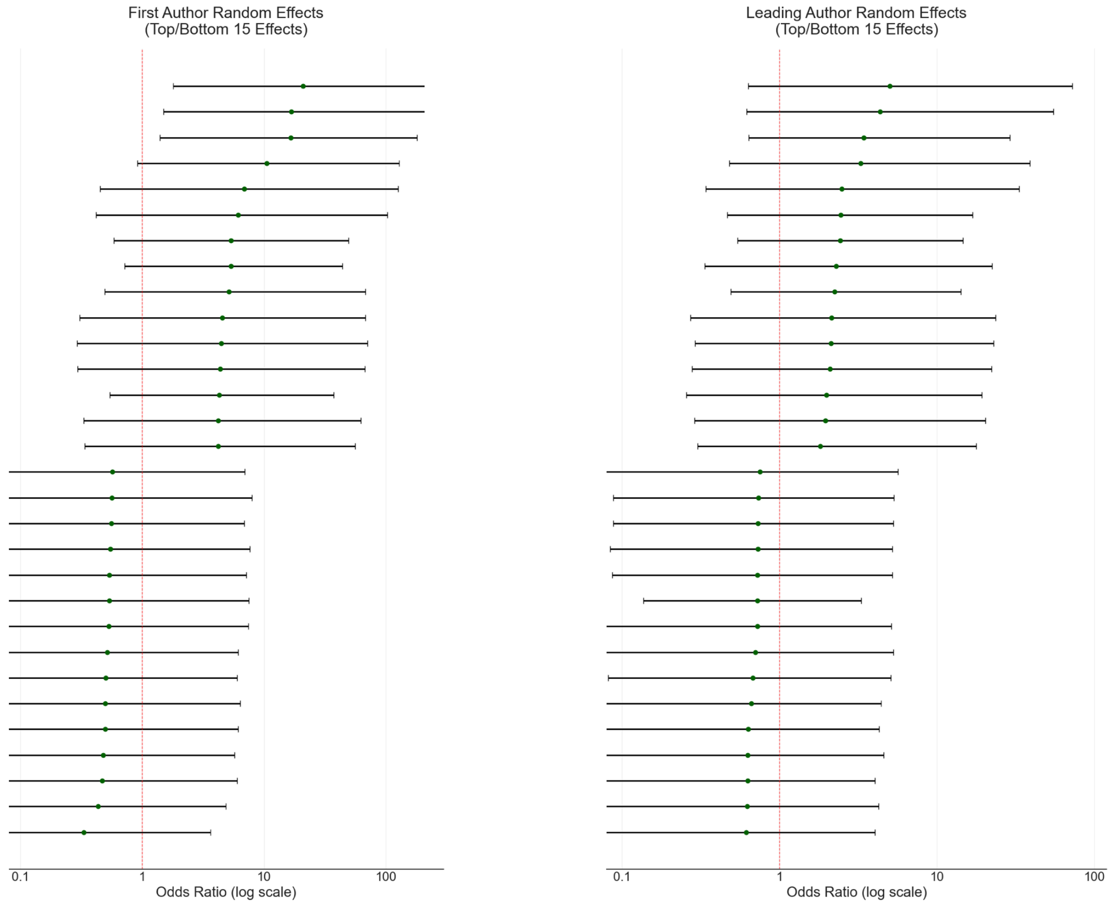


*Figure S2. Top and bottom 15 random effect for First author (left) and Leading author (right)*

#### All multivariate model fixed-effect results

Fixed-effect results from the multivariable Bayesian mixed-effects logistic regression of challenged claims.

| Domain | Predictor | Posterior β (SD) | Adjusted OR (94% HDI) |
|---|---|---|---|
| Journal impact factor | High-impact vs low-impact journal | 0.27 (0.41) | 1.31 (0.60–2.88) |
| | Trophy vs low-impact journal | 0.56 (0.56) | 1.75 (0.59–4.91) |
| Shanghai university rank | Ranked 101+ vs not ranked | 0.48 (0.53) | 1.61 (0.62–4.50) |

| Domain | Predictor | Posterior β (SD) | Adjusted OR (94% HDI) |
|---|---|---|---|
| | Ranked 51–100 vs not ranked | 0.14 (0.64) | 1.15 (0.34–3.81) |
| | Top 50 vs not ranked | 1.30 (0.48) | **3.68 (1.50–9.28)** |
| Publication year | B-spline basis 1 | 0.14 (0.99) | 1.15 (0.18–7.40) |
| | B-spline basis 2 | 0.25 (0.92) | 1.28 (0.23–6.99) |
| | B-spline basis 3 | -0.42 (0.85) | 0.66 (0.13–3.26) |
| First author | Female vs male | -0.06 (0.43) | 0.94 (0.41–2.11) |
| | Postdoctoral researcher vs PhD student | 0.15 (0.46) | 1.16 (0.49–2.77) |
| Leading author | Prior first-authorship in Drosophila immunity: yes vs no | -0.93 (0.61) | 0.39 (0.12–1.18) |
| | Original first-/last-author cohort vs no prior first-authorship | 0.30 (0.67) | 1.36 (0.38–4.69) |
| | Female vs male | 0.03 (0.56) | 1.03 (0.35–2.82) |
| | Junior vs senior PI | 0.27 (0.43) | 1.31 (0.58–2.94) |
| | Continuity vs exploratory research style | -0.12 (0.50) | 0.89 (0.35–2.33) |
| | First paper before 1995: yes vs no | -0.72 (0.71) | 0.49 (0.13–1.86) |

#### 3. Multivariate model using Unchallenged as outcome

We fit the multivariate model above using unchallenged as outcome. The methods are similar and the convergence and diagnosis checks are good, and not reported here for brevity. We observed the Odd-ratios shown in figure S3, and we note the significant effect, where author with a continuity style, have lower odds of having their claim remain unchallenged.

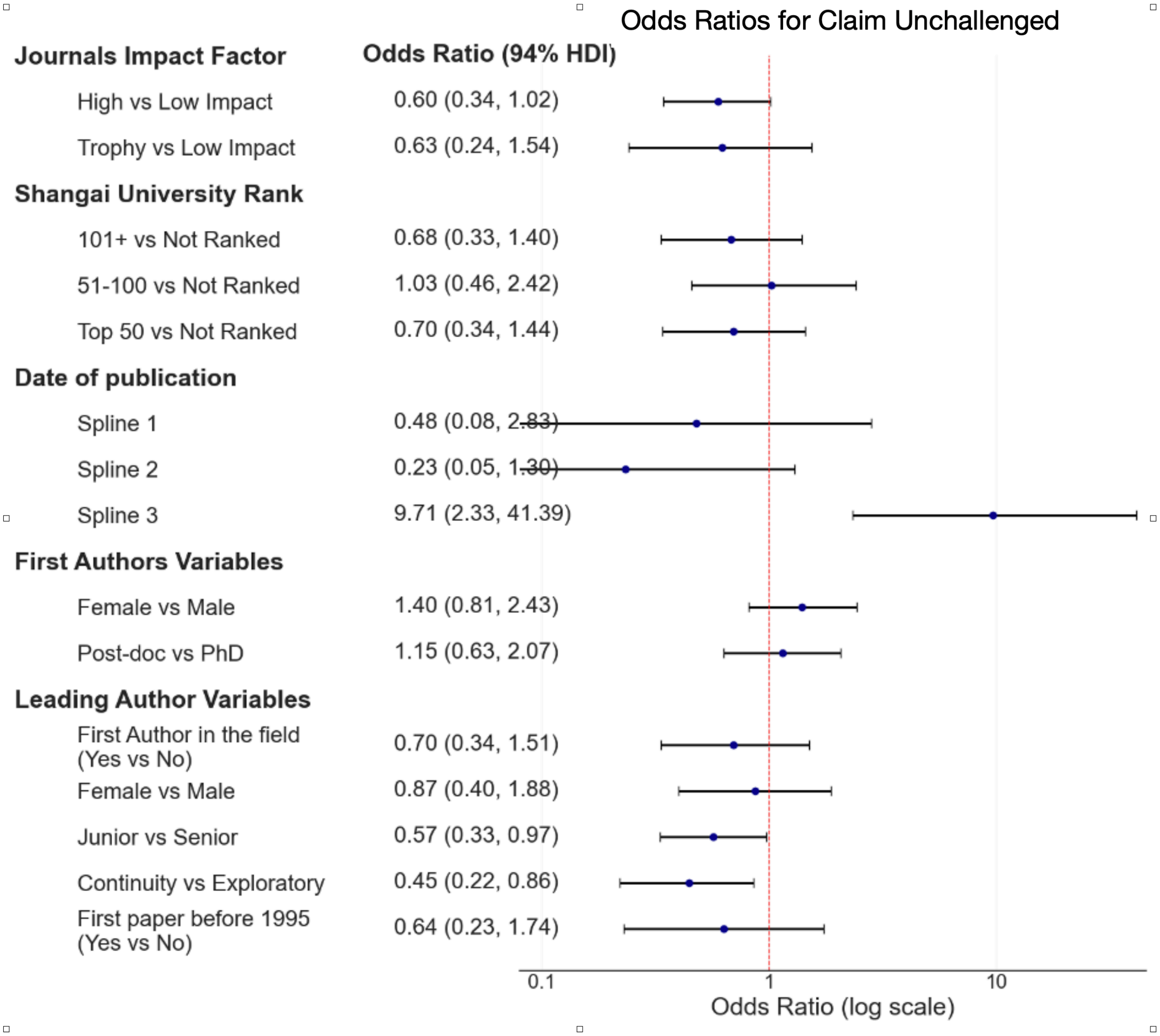


*Figure S3. Odd ratios for claim being unchallenged*

#### 4. Multivariate model excluding unchallenged claims

We fit the multivariate model from the main text while excluding unchallenged claims from the analysis. The methods are similar and the convergence and diagnosis checks are good and not reported here for brevity. The model simultaneously adjusts for author demographics, laboratory attributes and journal or institutional prestige and predicts a binary outcome for each claim: challenged or not challenged (including Verified, Partially Verified, Mixed). The re-analysis included 659 claims, of which 60 (9.1%) were challenged. The non-challenged reference group comprised of Verified (n=525), Partially Verified (n=64), Mixed (n=10). The Adjusted odds ratios and 94% highest-density intervals are reported in Figure S4. The results are very similar to the analysis with the unchallenged claims in the main text, with only university ranking (Top 50) being a significant predictor (OR 3.34 (1.28–9.01) vs the main text OR 3.68 (94% HDI 1.50–9.28). The other variable with the biggest changes shows only modest differences:

- high-impact journal OR 1.12 (vs 1.31)

- trophy journal OR 1.47 (vs 1.75)
- post-doc OR 1.44 (vs 1.16)

and their intervals still include 1. This model answers a different question from the main text model. The latter estimates the probability that a claim has been shown to be irreplicable to date, which includes the probability of being tested. The present model estimates the probability of failure among claims that were tested. Because being followed up is not random and depends on journal tier and institutional ranking (Figure S3), neither model is free of selection bias, and they should be interpreted together.

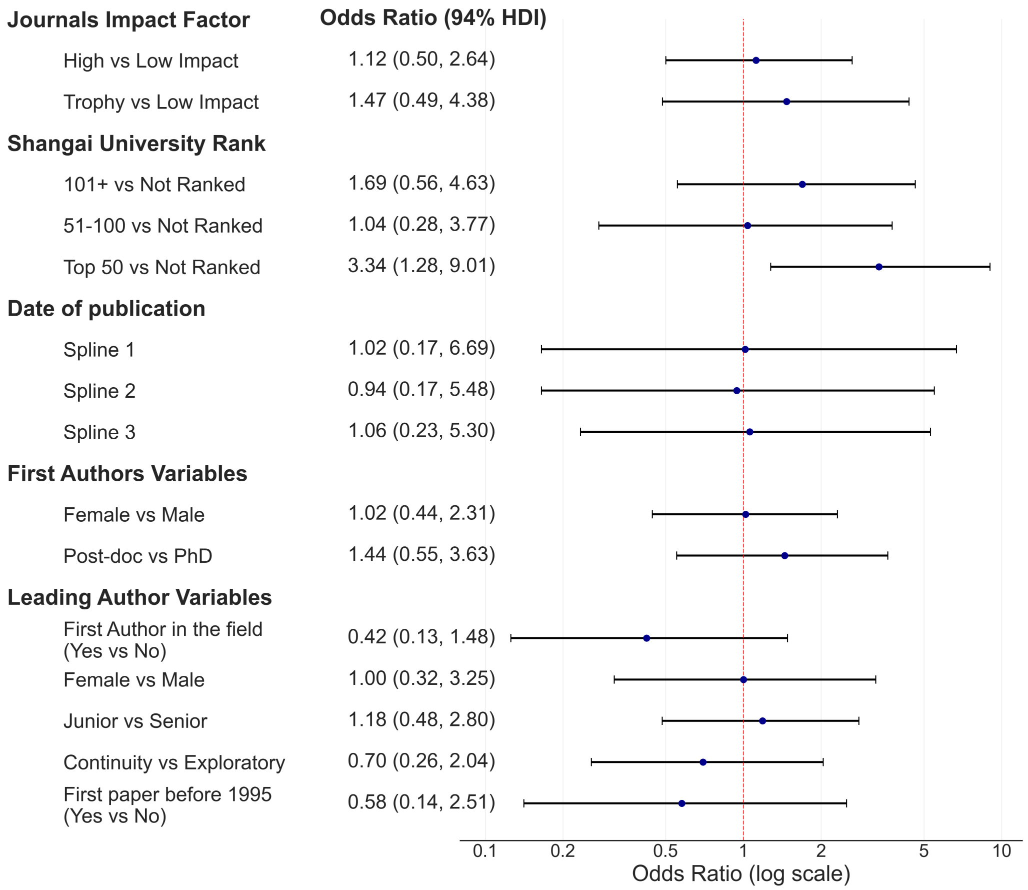


*Figure S4. Odd ratio for a model excluding unchallenged claims.*

#### 5. Multivariate model with continuous impact factor

As a sensitivity analysis, we refitted the multivariable hierarchical logistic model by replacing journal-impact-factor categories with $\log_2$-transformed continuous impact factor, while retaining all other covariates and random intercepts from the primary analysis. University ranking was retained as a categorical variable because the 2010 Shanghai Academic Ranking reports individual ranks only for institutions ranked 1–100 and otherwise reports rank bands (101–150,

151–200, 201–300, 301–400, and 401–500); moreover, 441 of 1,006 claims (43.8%) were from unranked institutions, which constitute a distinct group.

Each doubling of journal impact factor was associated with 1.51-fold higher odds that a claim was challenged (OR 1.51, 94% HDI 1.13–2.08). At the mean impact factors of the low-impact and trophy-journal tiers (4.4 and 61, respectively; approximately 3.8 doublings), the continuous model implies a trophy-versus-low-impact odds ratio of approximately 4.8. By contrast, the primary model directly estimated the trophy-versus-low-impact odds ratio as 1.75 (94% HDI 0.59–4.91). Although this wide interval includes 4.8, the point estimates differ between the two models. We interpret the larger implied trophy-journal effect in the continuous model cautiously because it assumes that the log-linear slope estimated predominantly from the many journals with lower impact factors (around 2–12) applies unchanged to the sparsely represented trophy journals (62 claims across *Science*, *Nature*, and *Cell*). We therefore retained the categorical journal analysis as the primary analysis because it directly estimates the trophy-journal comparison using claims published in those journals. Estimates for all other covariates were broadly similar to those in the primary model. In particular, the association with Top 50 university affiliation remained evident (OR 4.04, 94% HDI 1.57–10.08, compared with OR 3.68, 94% HDI 1.50–9.28 in the primary model).

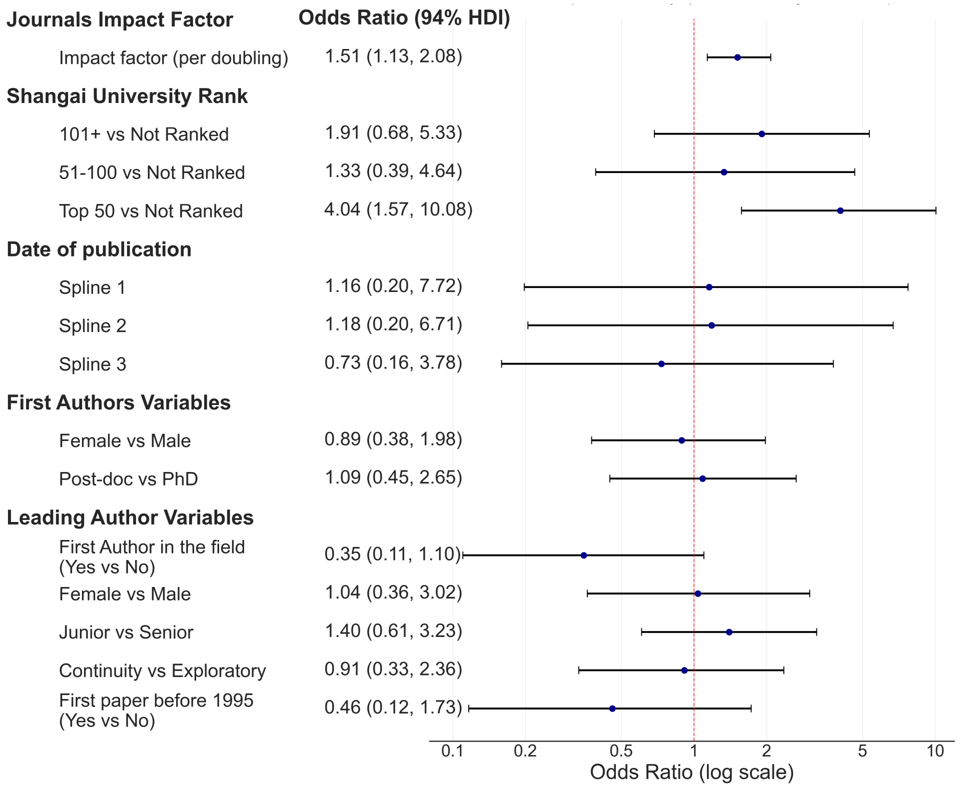


*Figure S5. Odd ratios for a model with impact factor modelled as a log-continuous predictor*

#### 6. Multivariate model with article level claims

Two claims in the same paper might not be independent, see Figure S6, so we might want to incorporate an article random effect into the primary Bayesian formulation.

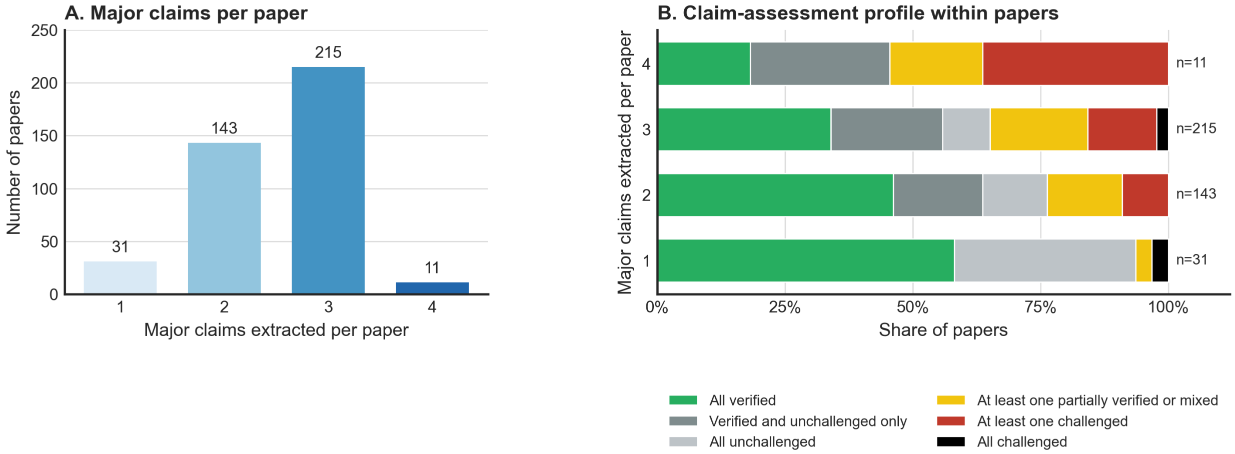


*Figure S6: A. Repartition of the claims in the articles: of the 400 papers, 215 contains 3 claims, and eleven contains 4 claims. B. Profile of the different articles' claim assessment.*

However, because the low number of claims per article (one to four) and only 43 articles (12.5%) contained at least one challenged claim, we could not manage to fit this model properly (we observed MCMC divergences). As a sensitivity analysis, we repeated the analysis with the article as the unit. Each of 345 articles with complete covariate data (representing 869 major claims, 60 of the 69 challenged claims in the full dataset; the remaining 9 fell in articles excluded for missing covariates) contributed as outcome. The outcome for an article was the proportion of its major claims classified as challenged (n_challenged/n_total_article_claims). The model was a (frequentist) binomial-link fractional-response generalized linear model with HC3 standard errors (the intervals reported here are 94% confidence intervals rather than posterior credible intervals), using the package statsmodels (Seabold and Perktold, 2010).

Each article contributed equally regardless of its number of claims. The Top 50 institutional effect remained significant under both analyses (OR 5.15, 94% CI 2.18–12.15; versus a claim-level OR 3.68, 94% HDI 1.50–9.28). In this new model Trophy journals also had higher odds of challenged claims than low-impact journals (OR 2.90, 94% CI 1.24–6.78 vs a claim-level OR 1.75, 94% HDI 0.59–4.91). This impact was not significant in the primary model but became significant in using article-level analysis. Other estimates are consistent in direction and magnitude with the primary claim-level model. Model diagnostics were satisfactory (Pearson dispersion = 0.49, design-matrix condition number = 18.9, no evidence of separation)

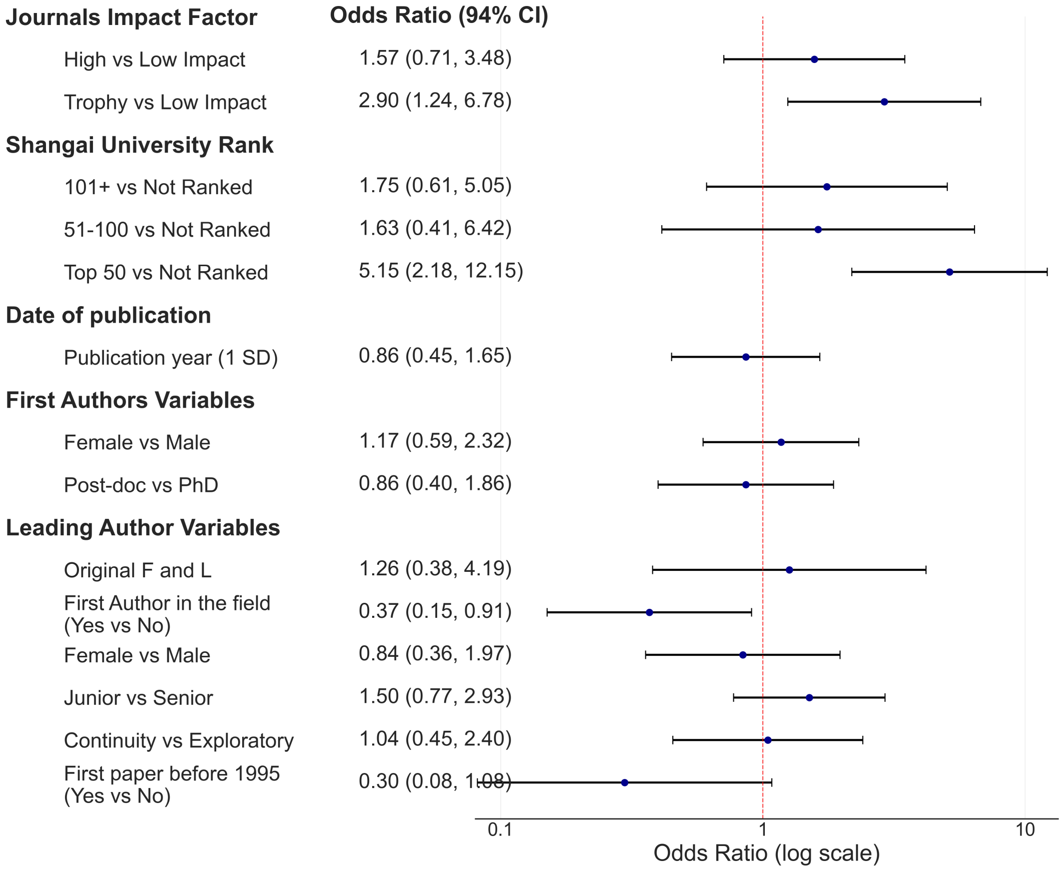


*Figure S7: odd-ratios for a model with article level claims.*

#### 7. Multivariate model using pre-experimental classifications

As a sensitivity analysis, we refitted the multivariable hierarchical logistic model using only the claim classifications available before experimental validation by the ReproSci project, as the choice of which claims to further validate could have been influenced by covariate (impact factor, university status). The 45 claims tested prospectively were therefore restored to their original "Unchallenged" classification, while all covariates and author-level random intercepts were retained from the primary analysis. The complete-case analysis included 869 claims, of which 28 (3.2%) were challenged. No predictor had a 94% highest-density interval excluding 1, including publication in trophy journals (OR 1.11, 94% HDI 0.33–3.77) and affiliation with a Top-50 institution (OR 1.44, 94% HDI 0.48–3.94). Thus, the associations observed in the pooled analysis were not evident when using classifications based exclusively on the retrospective literature review, although estimates were imprecise because of the small number of challenged claims.

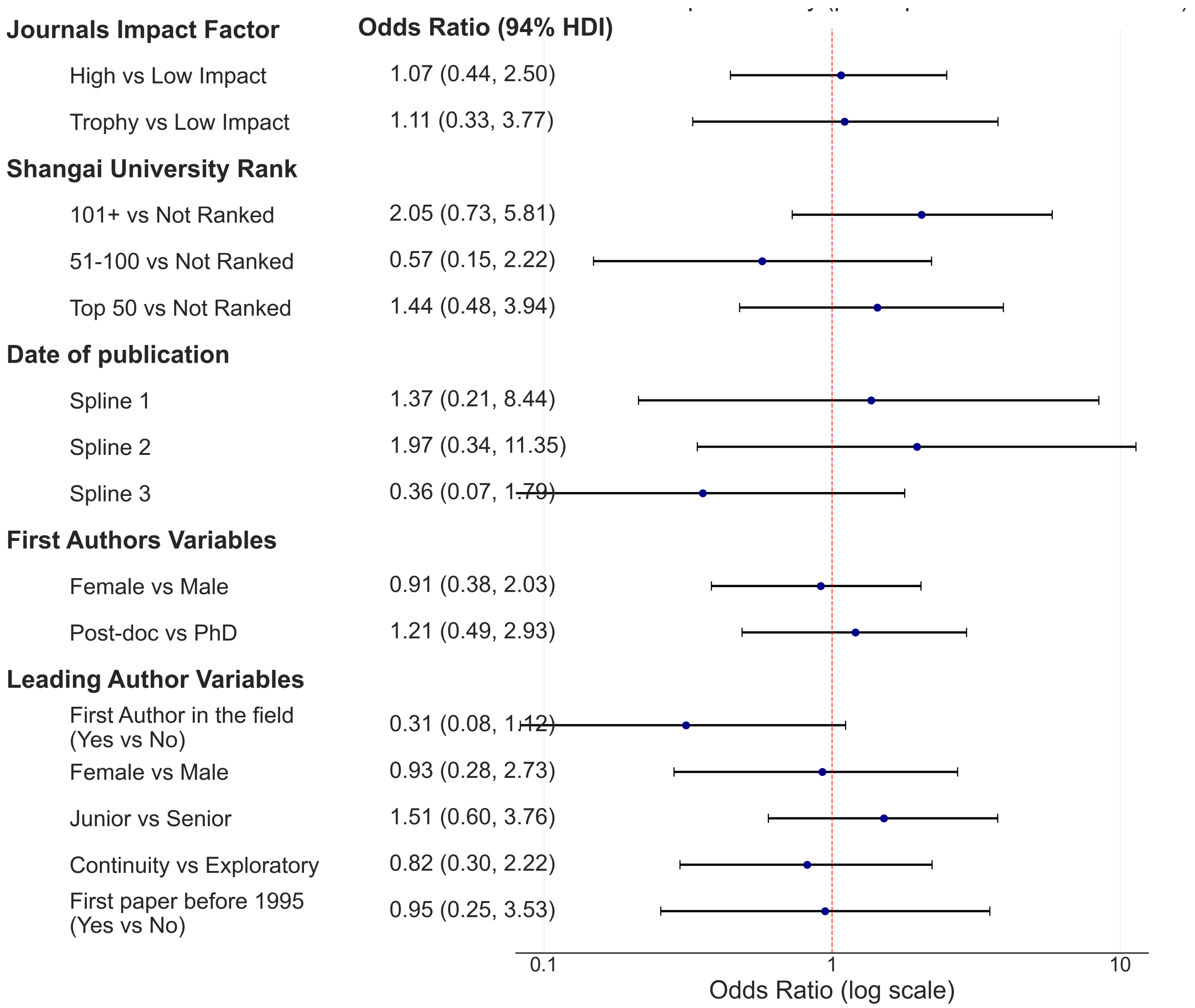


*Figure* ***Erreur ! Utilisez l'onglet Accueil pour appliquer 0 au texte que vous souhaitez faire apparaître ici.****:multivariate model using claims classification before validation by the ReproSci project*

### 8. Categorical comparison tables

We present table of the odd-rations and values for the categorical variables shown in the main text, for first and last-author.

#### First author

| *Variable* | Group 1 | Group 2 | Group1 Challenged % (95% CI) | Group2 Challenged % (95% CI) | Odds Ratio | OR 95% CI | Fisher p-value | Significance |
|---|---|---|---|---|---|---|---|---|
| *First Author Sex* | Male | Female | 7.3% (5.4–9.8%) | 6.7% (4.7–9.5%) | 1.09 | 0.66-1.78 | 0.802 | not significantly associated with |
| *First Author Become a PI* | TRUE | FALSE | 9.0% (6.6–12.2%) | 5.3% (3.8–7.5%) | 1.75 | 1.06-2.90 | 0.028 | significantly associated with |

#### Leading author

| Variable | Group1 | Group2 | Group1 Challenged % (95% CI) | Group2 Challenged % (95% CI) | Odds Ratio | OR 95% CI | Fisher p-value | Significance |
|---|---|---|---|---|---|---|---|---|
| *Trained in Historical laboratory (Comparison restricted to claims published after 1995)* | FALSE | TRUE | 8.7% (6.6–11.2%) | 5.0% (2.9–8.5%) | 1.82 | 0.95-3.47 | 0.082 | not significantly associated with |
| *Continuity* | FALSE | TRUE | 8.2% (5.8–11.5%) | 6.1% (4.5–8.2%) | 1.38 | 0.84-2.26 | 0.243 | not significantly associated with |
| *Leading Author Sex* | Male | Female | 7.0% (5.5–9.0%) | 5.8% (3.1–10.7%) | 1.22 | 0.59-2.51 | 0.729 | not significantly associated with |
| *Junior Senior* | Junior PI | Senior PI | 7.2% (5.1–10.2%) | 6.5% (4.8–8.8%) | 1.12 | 0.68-1.83 | 0.703 | not significantly associated with |
| *Was first author* | FALSE | TRUE | 8.2% (6.1–10.9%) | 3.5% (1.8–6.4%) | 2.48 | 1.19-5.18 | 0.013 | significantly associated with |

### 9. Permutation analysis of the Gini index

We compared the observed Gini coefficients with two permutation analyses, each based on 1,000,000 permutations. First, we randomly reassigned the challenged labels across individual claims while keeping every claim attached to its original author. This preserved the number of claims contributed by each author. Second, we kept the challenged-claim pattern of each paper together and reassigned these patterns among papers containing the same number of claims. This additionally accounted for the possibility that claims from the same paper were challenged because of a shared problem. For each permutation, we recalculated the Gini coefficient across authors and calculated the p-value as the proportion of simulated coefficients that equaled or exceeded the observed coefficient.

| Author role | Permutation | Observed Gini | Mean null Gini | 95% interval | p |
|---|---|---|---|---|---|
| First author | Individual claims | 0.881 | 0.808 | 0.781–0.834 | 0.000001 |
| First author | Papers kept together | 0.881 | 0.866 | 0.856–0.879 | 0.015 |
| Leading author | Individual claims | 0.856 | 0.784 | 0.733–0.831 | 0.00081 |
| Leading author | Papers kept together | 0.856 | 0.834 | 0.798–0.870 | 0.127 |

### 10. Other comparisons

#### Journal category proportions and comparisons for challenged as outcome

*Challenged proportions by journal category*:

- Low Impact: 5.8% (95% CI 4.3–7.9%) of Low Impact claims were challenged.
- High Impact: 7.8% (95% CI 5.3–11.3%) of High Impact claims were challenged.
- Trophy Journals: 12.9% (95% CI 6.7–23.4%) of Trophy Journals claims were challenged.

*Statistical comparisons (using Low-Impact as baseline):*

- High-impact vs low-impact journal category high impact vs low impact was not significantly associated with claim replicability (p = 0.26); 7.8% (95% CI 5.3–11.3%) vs 5.8% (95% CI 4.3–7.9%) of claims were challenged for High Impact vs Low Impact, respectively. Odds Ratio = 1.36 (95% CI 0.80–2.32).
- Trophy vs low-impact journal category trophy journals vs low impact was not significantly associated with claim replicability (p = 0.05); 12.9% (95% CI 6.7–23.4%) vs 5.8% (95% CI 4.3–7.9%) of claims were challenged for Trophy Journals vs Low Impact, respectively. Odds Ratio = 2.39 (95% CI 1.06–5.40).
- Trophy vs high-impact journal category trophy journals vs high impact was not significantly associated with claim replicability (p = 0.21); 12.9% (95% CI 6.7–23.4%) vs 7.8% (95% CI 5.3–11.3%) of claims were challenged for Trophy Journals vs High Impact, respectively. Odds Ratio = 1.76 (95% CI 0.75–4.12).

#### Journal category proportions and comparisons for **unchallenged** as outcome

*Unchallenged proportions by journal category*:

- Low Impact: 29.3% (95% CI 25.9–32.9%) of Low Impact claims were unchallenged.
- High Impact: 13.9% (95% CI: 10.5%–18.2%) of High Impact claims were unchallenged.
- Trophy Journals: 17.7% (95% CI: 10.2%–29.0%) of Trophy Journals claims were unchallenged.

*Statistical comparisons (using Low-Impact as baseline):*

- High-impact vs low-impact journal category high impact vs low impact was significantly associated with claim replicability (p = 0.00); 29.3% (95% CI 25.9–32.9%) vs 13.9% (95% CI 10.5–18.2%) of claims were unchallenged for Low Impact vs High Impact, respectively. Odds Ratio = 2.56 (95% CI 1.78–3.69).
- Trophy vs low-impact journal category trophy journals vs low impact was not significantly associated with claim replicability (p = 0.06); 29.3% (95% CI 25.9–32.9%) vs 17.7% (95% CI 10.2–29.0%) of claims were unchallenged for Low Impact vs Trophy Journals, respectively. Odds Ratio = 1.92 (95% CI 0.98–3.77).
- Trophy vs high-impact journal category trophy journals vs high impact was not significantly associated with claim replicability (p = 0.43); 17.7% (95% CI 10.2–29.0%) vs 13.9% (95% CI 10.5–18.2%) of claims were unchallenged for Trophy Journals vs High Impact, respectively. Odds Ratio = 1.33 (95% CI 0.65–2.76).

#### University Ranking category proportions and comparisons for challenged as outcome

##### *Challenged proportions by university ranking:*

- Top 50: 15.8% (95% CI: 11.4%–21.6%) of Top 50 university claims were challenged.
- 51-100: 6.0% (95% CI: 3.1%–11.4%) of 51-100 university claims were challenged.
- 101+: 5.5% (95% CI: 3.2%–9.2%) of 101+ university claims were challenged.
- Not Ranked: 3.9% (95% CI: 2.4%–6.1%) of Not Ranked university claims were challenged.

##### *University ranking statistical comparisons*

- Top 50 vs not ranked university top 50 vs not ranked was significantly associated with claim replicability (p = 0.00); 15.8% (95% CI 11.4–21.6%) vs 3.9% (95% CI 2.4–6.1%) of claims were challenged for Top 50 vs Not Ranked, respectively. Odds Ratio = 4.69 (95% CI 2.53–8.70).
- 51-100 ranked vs not ranked university 51-100 vs not ranked was not significantly associated with claim replicability (p = 0.33); 6.0% (95% CI 3.1–11.4%) vs 3.9% (95% CI 2.4–6.1%) of claims were challenged for 51-100 vs Not Ranked, respectively. Odds Ratio = 1.60 (95% CI 0.67–3.79).
- 101+ ranked vs not ranked university 101+ vs not ranked was not significantly associated with claim replicability (p = 0.33); 5.5% (95% CI 3.2–9.2%) vs 3.9% (95% CI 2.4–6.1%) of claims were challenged for 101+ vs Not Ranked, respectively. Odds Ratio = 1.45 (95% CI 0.69–3.05).

#### Time Period proportions and comparisons for challenged as outcome

##### *Challenged proportions by time period:*

- ≤1991: 0.0% (95% CI: 0.0%–11.7%) of ≤1991 claims were challenged.
- 1992-1996: 4.6% (95% CI: 1.8%–11.2%) of 1992-1996 claims were challenged.
- 1997-2001: 7.7% (95% CI: 4.5%–13.0%) of 1997-2001 claims were challenged.
- 2002-2006: 8.3% (95% CI: 5.9%–11.5%) of 2002-2006 claims were challenged.
- 2007-2011: 6.1% (95% CI: 4.1%–9.1%) of 2007-2011 claims were challenged.

##### *Time period statistical comparisons (pairwise)*

- 1992-1996 vs ≤1991 was not significantly associated with claim replicability (p = 0.57); 4.6% (95% CI 1.8–11.2%) vs 0.0% (95% CI 0.0–11.7%) of claims were challenged for 1992-1996 vs ≤1991, respectively. Odds Ratio = 2.80 (95% CI 0.14–54.49).
- 1997-2001 vs 1992-1996 was not significantly associated with claim replicability (p = 0.43); 7.7% (95% CI 4.5–13.0%) vs 4.6% (95% CI 1.8–11.2%) of claims were

challenged for 1997-2001 vs 1992-1996, respectively.  Odds Ratio = 1.74 (95% CI 0.54–5.57).

- 2002-2006 vs 1997-2001 was not significantly associated with claim replicability (p = 1.00);  8.3% (95% CI 5.9–11.5%) vs 7.7% (95% CI 4.5–13.0%) of claims were challenged for 2002-2006 vs 1997-2001, respectively. Odds Ratio = 1.08 (95% CI 0.54–2.16).
- 2007-2011 vs 2002-2006 was not significantly associated with claim replicability (p = 0.26); 8.3% (95% CI 5.9–11.5%) vs 6.1% (95% CI 4.1–9.1%) of claims were challenged for 2002-2006 vs 2007-2011, respectively.  Odds Ratio = 1.39 (95% CI 0.79–2.45).
- 2002-2006 vs ≤1991 was not significantly associated with claim replicability (p = 0.15); 8.3% (95% CI 5.9–11.5%) vs 0.0% (95% CI 0.0–11.7%) of claims were challenged for 2002-2006 vs ≤1991, respectively.  Odds Ratio = 5.24 (95% CI 0.31–87.91).

#### 11. Table of journals with 2022 impact factor, and claim assessment

| journal_name | impact_factor | Challenged | Mixed | Partially Verified | Unchallenged | Verified | Total |
|---|---|---|---|---|---|---|---|
| **Nature** | 64.8 | 3 | 0 | 0 | 7 | 10 | 20 |
| **Cell** | 64.5 | 2 | 0 | 2 | 1 | 10 | 15 |
| **Science (New York, N.Y.)** | 56.9 | 3 | 1 | 3 | 3 | 17 | 27 |
| **Immunity** | 32.4 | 2 | 0 | 1 | 0 | 21 | 24 |
| **Nature immunology** | 30.5 | 4 | 1 | 1 | 3 | 19 | 28 |
| **Cell host & microbe** | 30.3 | 1 | 0 | 1 | 2 | 7 | 11 |
| **Molecular cell** | 16 | 0 | 0 | 1 | 2 | 5 | 8 |
| **Nucleic acids research** | 14.9 | 0 | 0 | 0 | 3 | 8 | 11 |
| **Nature protocols** | 14.8 | 0 | 0 | 0 | 0 | 1 | 1 |
| **Developmental cell** | 11.8 | 1 | 0 | 1 | 7 | 15 | 24 |
| **The EMBO journal** | 11.4 | 7 | 0 | 5 | 3 | 41 | 56 |
| **Proceedings of the National Academy of Sciences of the United States of America** | 11.1 | 6 | 2 | 11 | 23 | 77 | 119 |
| **Genes & development** | 10.5 | 3 | 0 | 1 | 0 | 21 | 25 |
| **National Cancer Institute monograph** | 10.3 | 0 | 0 | 0 | 0 | 2 | 2 |
| **PLoS biology** | 9.8 | 5 | 0 | 1 | 10 | 3 | 19 |
| **Experientia** | 9.261 | 0 | 0 | 0 | 1 | 0 | 1 |
| **Current biology : CB** | 9.2 | 3 | 0 | 2 | 12 | 19 | 36 |
| **EMBO reports** | 9.071 | 2 | 0 | 3 | 8 | 18 | 31 |
| **The Journal of biophysical and biochemical cytology** | 7.8 | 0 | 0 | 1 | 0 | 1 | 2 |
| **The Journal of cell biology** | 7.8 | 1 | 0 | 0 | 0 | 0 | 1 |
| **Aging cell** | 7.8 | 1 | 0 | 0 | 1 | 4 | 6 |
| **Molecular & cellular proteomics : MCP** | 7 | 0 | 0 | 1 | 0 | 2 | 3 |
| **Journal of experimental botany** | 6.9 | 0 | 0 | 0 | 0 | 2 | 2 |
| **PLoS pathogens** | 6.7 | 4 | 0 | 2 | 19 | 15 | 40 |
| **Microbes and infection** | 5.8 | 0 | 0 | 1 | 3 | 3 | 7 |
| **Journal of the American Aging Association** | 5.6 | 0 | 0 | 1 | 0 | 2 | 3 |
| **Journal of molecular biology** | 5.6 | 0 | 0 | 0 | 0 | 4 | 4 |
| **European journal of biochemistry** | 5.4 | 1 | 0 | 1 | 0 | 10 | 12 |
| **European journal of immunology** | 5.4 | 0 | 0 | 0 | 3 | 0 | 3 |
| **The FEBS journal** | 5.4 | 0 | 0 | 0 | 1 | 2 | 3 |
| **Journal of innate immunity** | 5.3 | 3 | 0 | 1 | 0 | 10 | 14 |
| **Journal of endotoxin research** | 5.3 | 1 | 0 | 0 | 0 | 4 | 5 |
| **Molecular and cellular biology** | 5.3 | 0 | 2 | 0 | 7 | 12 | 21 |
| **Virulence** | 5.2 | 0 | 0 | 0 | 0 | 1 | 1 |
| **Molecular ecology** | 4.9 | 0 | 0 | 1 | 1 | 0 | 2 |
| **The Journal of biological chemistry** | 4.8 | 8 | 2 | 7 | 15 | 41 | 73 |

| | | | | | | | |
|---|---|---|---|---|---|---|---|
| **FASEB journal : official publication of the Federation of American Societies for Experimental Biology** | 4.8 | 0 | 0 | 1 | 1 | 4 | 6 |
| **Cellular signalling** | 4.8 | 0 | 0 | 1 | 1 | 4 | 6 |
| **Proceedings Biological Sciences** | 4.7 | 0 | 0 | 4 | 1 | 1 | 6 |
| **Development (Cambridge, England)** | 4.6 | 0 | 1 | 0 | 1 | 1 | 3 |
| **PLoS genetics** | 4.5 | 0 | 0 | 0 | 2 | 1 | 3 |
| **Journal of immunology (Baltimore, Md. : 1950)** | 4.4 | 1 | 1 | 0 | 2 | 9 | 13 |
| **BMC genomics** | 4.4 | 1 | 0 | 0 | 4 | 2 | 7 |
| **Immunology letters** | 4.4 | 0 | 0 | 1 | 0 | 1 | 2 |
| **Cellular immunology** | 4.3 | 1 | 0 | 0 | 0 | 1 | 2 |
| **Disease models & mechanisms** | 4.3 | 0 | 0 | 0 | 0 | 3 | 3 |
| **The Biochemical journal** | 4.1 | 0 | 0 | 0 | 2 | 1 | 3 |
| **Journal of cellular biochemistry** | 4 | 0 | 0 | 0 | 3 | 3 | 6 |
| **Journal of cell science** | 4 | 0 | 0 | 2 | 11 | 5 | 18 |
| **Experimental gerontology** | 3.9 | 0 | 0 | 0 | 0 | 2 | 2 |
| **Cytokine** | 3.8 | 0 | 0 | 0 | 1 | 0 | 1 |
| **Insect biochemistry and molecular biology** | 3.8 | 1 | 1 | 1 | 2 | 16 | 21 |
| **Molecules and cells** | 3.8 | 0 | 0 | 0 | 1 | 4 | 5 |
| **Experimental cell research** | 3.7 | 0 | 0 | 0 | 1 | 2 | 3 |
| **PloS one** | 3.7 | 0 | 0 | 0 | 9 | 4 | 13 |
| **Advances in experimental medicine and biology** | 3.65 | 0 | 0 | 0 | 0 | 1 | 1 |
| **Molecular immunology** | 3.6 | 1 | 0 | 2 | 4 | 6 | 13 |
| **Gene** | 3.5 | 0 | 0 | 1 | 1 | 3 | 5 |
| **FEBS letters** | 3.5 | 0 | 0 | 0 | 1 | 2 | 3 |
| **Cellular microbiology** | 3.4 | 0 | 0 | 0 | 3 | 12 | 15 |
| **Differentiation; research in biological diversity** | 3.39 | 0 | 0 | 0 | 1 | 1 | 2 |
| **Genetics** | 3.3 | 0 | 0 | 3 | 2 | 16 | 21 |
| **Journal of bacteriology** | 3.2 | 0 | 0 | 0 | 1 | 2 | 3 |
| **Developmental biology** | 3.148 | 0 | 1 | 2 | 1 | 15 | 19 |
| **Molecular & general genetics : MGG** | 3.1 | 0 | 0 | 0 | 2 | 7 | 9 |
| **Biochemical and biophysical research communications** | 3.1 | 1 | 0 | 1 | 4 | 20 | 26 |
| **Infection and immunity** | 3.1 | 0 | 0 | 1 | 5 | 3 | 9 |
| **Biochimica et biophysica acta** | 3 | 0 | 0 | 0 | 4 | 0 | 4 |
| **Proteins** | 3 | 1 | 0 | 0 | 0 | 1 | 2 |
| **Eukaryotic cell** | 2.992 | 0 | 0 | 0 | 5 | 0 | 5 |
| **Developmental and comparative immunology** | 2.9 | 0 | 0 | 0 | 7 | 14 | 21 |
| **The Journal of experimental biology** | 2.8 | 0 | 0 | 0 | 1 | 2 | 3 |
| **Molecular biology reports** | 2.8 | 0 | 0 | 0 | 0 | 1 | 1 |
| **Insect molecular biology** | 2.6 | 0 | 0 | 0 | 4 | 6 | 10 |
| **Mechanisms of development** | 2.6 | 0 | 0 | 0 | 3 | 3 | 6 |
| **International journal of immunogenetics** | 2.2 | 0 | 0 | 0 | 3 | 0 | 3 |
| **Journal of insect physiology** | 2.2 | 0 | 0 | 1 | 1 | 6 | 8 |
| **Genes to cells : devoted to molecular & cellular mechanisms** | 2.1 | 0 | 0 | 2 | 7 | 2 | 11 |
| **Journal of evolutionary biology** | 2.1 | 0 | 0 | 0 | 1 | 2 | 3 |
| **BMC developmental biology** | 1.978 | 0 | 0 | 0 | 0 | 2 | 2 |
| **Protein expression and purification** | 1.6 | 0 | 0 | 0 | 0 | 1 | 1 |
| **Journal of morphology** | 1.5 | 0 | 0 | 2 | 0 | 0 | 2 |
| **Gene expression patterns : GEP** | 1.2 | 0 | 0 | 1 | 1 | 0 | 2 |
| **Fly** | 1.2 | 0 | 0 | 0 | 1 | 5 | 6 |
| **Comptes rendus de l'Academie des sciences. Serie III, Sciences de la vie** | 0.8 | 1 | 0 | 0 | 0 | 2 | 3 |
| **Acta biologica Hungarica** | 0.585 | 0 | 0 | 0 | 0 | 2 | 2 |
| **Annales de parasitologie humaine et comparee** | 0.5 | 0 | 0 | 0 | 0 | 2 | 2 |

*Table S1. List of journals with impact factor and claim assessment*

#### 12. References

Abril-Pla O, Andreani V, Carroll C, Dong L, Fonnesbeck CJ, Kochurov M, Kumar R, Lao J, Luhmann CC, Martin OA, Osthege M, Vieira R, Wiecki T, Zinkov R. 2023. PyMC: a modern, and comprehensive probabilistic programming framework in Python. PeerJ Computer Science **9**:e1516. DOI: https://doi.org/10.7717/peerj-cs.1516

Capretto T, Piho C, Kumar R, Westfall J, Yarkoni T, Martin OA. 2022. Bambi: A Simple Interface for Fitting Bayesian Linear Models in Python. Journal of Statistical Software **103**:1–29. DOI: https://doi.org/10.18637/jss.v103.i15

Grinsztajn L, Semenova E, Margossian CC, Riou J. 2021. Bayesian workflow for disease transmission modeling in Stan. Statistics in Medicine **40**:6209–6234. DOI: https://doi.org/10.1002/sim.9164

Seabold S, Perktold J. 2010. Statsmodels: Econometric and Statistical Modeling with Python. . Presented at the Python in Science Conference. p. 92–96. DOI: https://doi.org/10.25080/Majora-92bf1922-011
